# The Mammalian Cytosolic Type 2 (R)-β-hydroxybutyrate Dehydrogenase (BDH2) is 4-oxo-L-proline Reductase (EC 1.1.1.104)

**DOI:** 10.1101/2021.02.23.432487

**Authors:** Sebastian P. Kwiatkowski, Maria Bozko, Michal Zarod, Apolonia Witecka, Adam K. Jagielski, Jakub Drozak

## Abstract

The early studies on chicken embryos revealed that exposition to 4-oxo-L-proline resulted in the explicit increase in 4-hydroxy-L-proline content in their tissues. In 1962, 4-oxo-L-proline reductase, an enzyme responsible for the reduction of 4-oxo-L-proline, was partially purified from rabbit kidneys and characterized biochemically, but only recently the molecular identity of the enzyme has been unveiled in our laboratory. The present investigation reports the purification, identification as well as biochemical characterization of 4-oxo-L-proline reductase. The enzyme was purified from rat kidneys about 280-fold. Following mass spectrometry analysis of the purified protein preparation, the mammalian cytosolic type 2 (R)-β-hydroxybutyrate dehydrogenase (BDH2) emerged as the only meaningful candidate for the reductase. Rat and human BDH2 were expressed in *E. coli*, purified, and shown to catalyze the reversible reduction of 4-oxo-L-proline to *cis*-4-hydroxy-L-proline, as confirmed by chromatographic and mass spectrometry analysis. Specificity studies carried out on both enzymes showed that 4-oxo-L-proline was the best substrate, particularly the human enzyme acted with 9400-fold higher catalytic efficiencies on 4-oxo-L-proline than on (R)-β-hydroxybutyrate. Finally, HEK293T cells efficiently metabolized 4-oxo-L-proline to *cis*-4-hydroxy-L-proline and simultaneously accumulated *trans*-4-hydroxy-L-proline in the culture medium, suggesting that 4-oxo-L-proline is most likely an inhibitor of *trans*-4-hydroxy-L-proline metabolism in human cells. We conclude that BDH2 is mammalian 4-oxo-L-proline reductase that converts 4-oxo-L-proline to *cis*-4-hydroxy-L-proline, and not to *trans*-4-hydroxy-L-proline as currently thought, and hypothesize that the enzyme may be considered as a potential source of *cis*-4-hydroxy-L-proline in mammalian tissues.

## Introduction

4-oxo-L-proline is a poorly studied derivative of L-proline (Fig. 1). Although the only known natural source of this compound is antibiotic X-type actinomycin produced by *Streptomyces antibioticus* (1), 4-oxo-L-proline has been occasionally detected in various biological samples, including extracts of human embryonic kidney 293 cells (HEK293T) (2) as well as the blood samples of type 2 diabetes patients treated with metformin, sulphonylurea or both drugs combined (3). Unfortunately, the exact source and metabolic routes of 4-oxo-L-proline *in vivo* have never been addressed appropriately and our knowledge of its physiological significance is still sparse. Interestingly, in the late 1950s, Mitoma and coworkers (4) showed that the administration of 4-oxo-L-proline to chick embryos increased the free 4-hydroxy-L-proline content in their tissues, suggesting that 4-oxo-L-proline might be enzymatically reduced *in vivo*. Later, 4-oxo-L-proline reductase (EC 1.1.1.104), a cytosolic enzyme converting 4-oxo-L-proline into 4-hydroxy-L-proline at the expense of NADH oxidation (*cf*. Fig. 1), was partially purified from rabbit kidney and characterized (5), but its molecular identity and biochemical properties have remained unknown so far.

**Figure 1.**
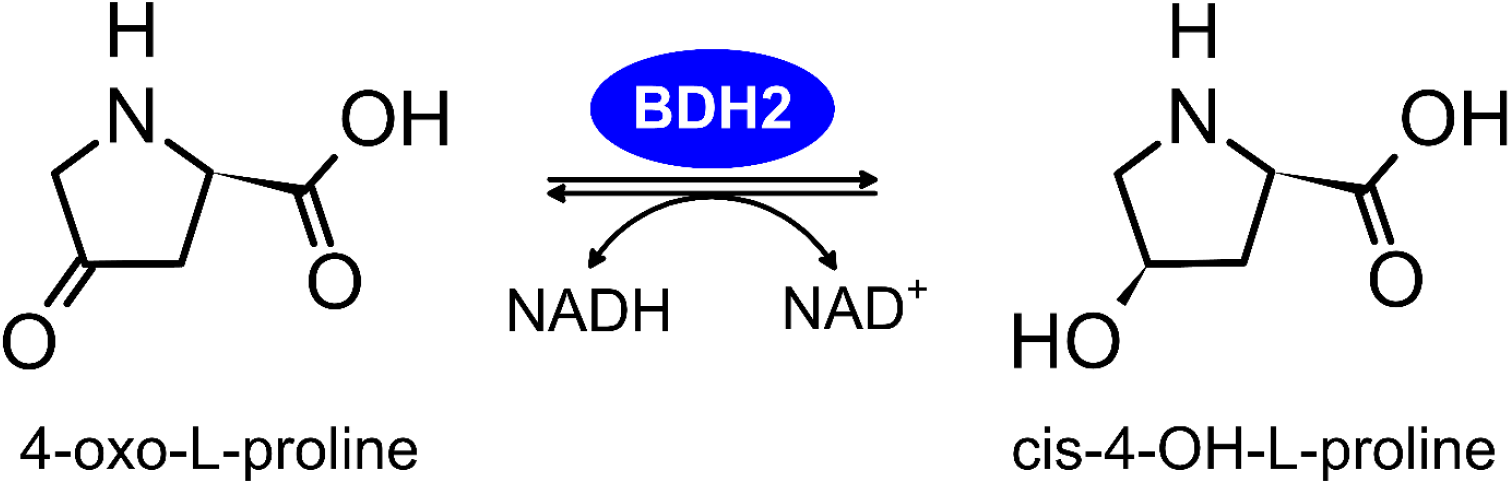
Scheme of the reaction catalyzed by BDH2 as a 4-oxo-L-proline reductase. BDH2 also acts as a *cis*-4-OH-L-proline dehydrogenase in the presence of the NAD^+^ *in vitro*. The biological relevance of the reverse reaction remains to be explored.

In the current work, we report the identification of mammalian 4-oxo-L-proline reductase (EC 1.1.1.104) as 3-hydroxybutyrate dehydrogenase type 2 (BDH2, DHRS6, Fig. 2), which was previously suggested to act as a cytosolic (R)-β-hydroxybutyrate dehydrogenase involved in ketone body utilization (6) or to catalyze the synthesis of 2,5-dihydroxybenzoic acid (2,5-DHBA, gentisic acid), a putative mammalian siderophore (7). We provide a biochemical characterization of this enzyme and show that it catalyzes the reversible conversion of 4-oxo-L-proline to *cis*-4-hydroxy-L-proline, with favorable substrate specificity and catalytic efficiency. BDH2 is therefore the first mammalian enzyme capable of yielding *cis*-4-hydroxy-L-proline, a metabolite thought to only be of exogenous origin. We also reveal that HEK293T cells efficiently metabolize 4-oxo-L-proline to *cis*-4-hydroxy-L-proline, indicating that the activity of BDH2 might be considered as a potential source of *cis*-4-hydroxy-L-proline in mammalian tissues. However, the physiological importance of these findings remains unclear.

**Figure 2.**
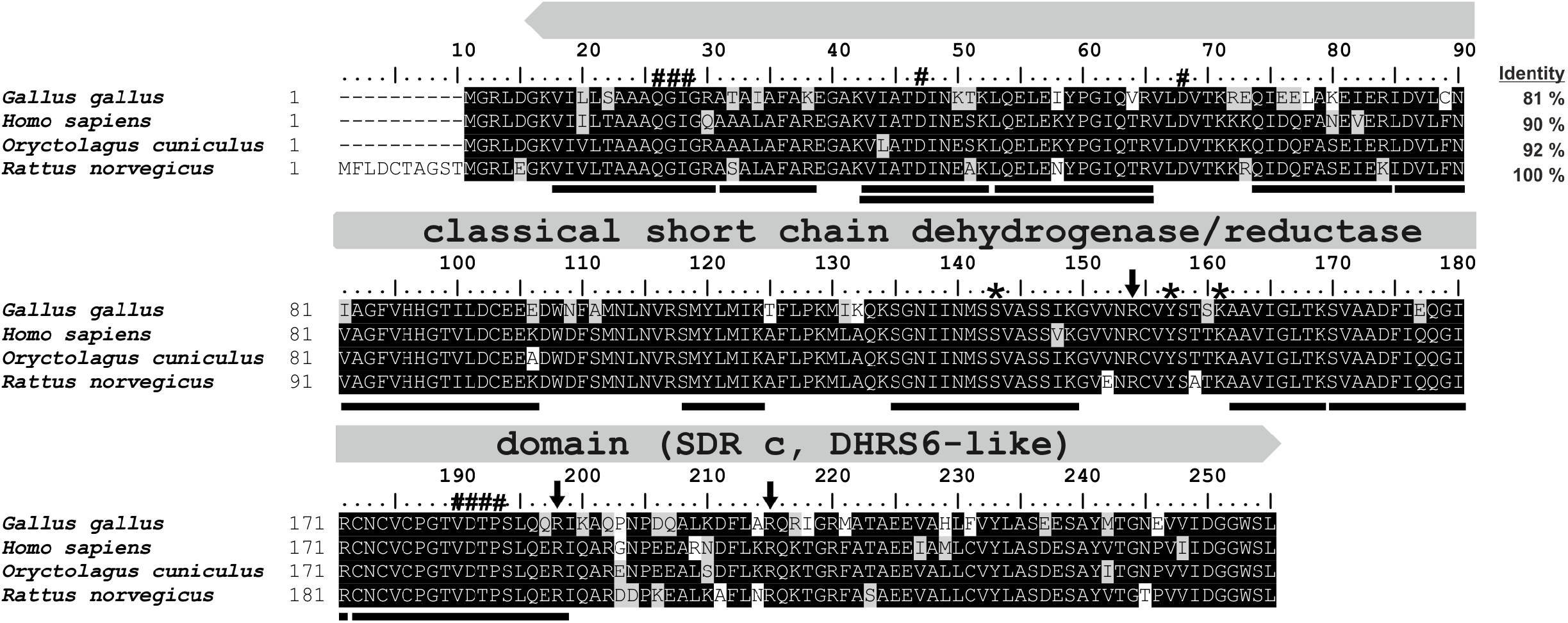
Amino acid sequence alignment of rat BDH2 protein with its orthologues. Sequences of rat (*Rattus norvegicus*, NP_001099943.1), human (*Homo sapiens*, NP_064524.3), rabbit (*Oryctolagus cuniculus*, XP_002717250.1), and chicken (*Gallus gallus*, XP_015141101.1) protein were obtained from the NCBI Protein database. Percentage of amino acid identities with rat BDH2 protein is given in the upper right. The characteristic one-domain architecture of the enzyme is indicated by the label above the alignment. Amino acid residues interacting with NAD(H) are indicated by hashes, whereas arginine residues coordinating the carboxylic group of a substrate are shown by arrows. Asterisks mark the key catalytic residues (6). The peptides identified by mass spectrometry in the protein purified from rat kidneys are underlined in the rat sequence, and several peptides covering similar though shorter sequences have been omitted. The level of residues conservation is indicated by black (100%) and gray (50% and more) background.

## Results

### Purification and identification of rat 4-oxo-L-proline reductase

Through the purification process, 4-oxo-L-proline reductase was assayed spectrophotometrically by measuring the rate of 4-oxo-L-proline reduction with concomitant oxidation of NADH to NAD^+^. The enzyme was purified from rat kidneys about 280-fold by a three-step procedure of column chromatography involving anion-exchange chromatography on Q Sepharose FF resin, affinity chromatography on HiScreen Blue FF column and gel filtration on Superdex 200 16/60 HiLoad column (Fig. 3). Only one enzyme species was present throughout the purification process as indicated by a single activity peak at each purification step. The yield of the purification was 25%. For details on the purification refer to Table 1.

**Table 1.** Purification of 4-oxo-L-proline reductase from rat kidneys.

| **Fraction** | **Volume** | **Total protein** | **Total activity** | **Specific activity** | **Purification** | **Yield** |
| --- | --- | --- | --- | --- | --- | --- |
|  | *ml* | *mg* | *$\mu\text{mol min}^{-1}$* | *$\mu\text{mol min}^{-1} \text{mg}^{-1}$* | *-fold* | *%* |
| 10% - 20% PEG | 120 | 2343.75 | 13.666 | 0.006 | 1 | 100 |
| Q Sepharose FF | 20 | 148.264 | 7.058 | 0.048 | 8 | 52 |
| HiScreen Blue FF | 12 | 16.734 | 3.055 | 0.183 | 31 | 22 |
| Superdex 200 16/60 HiLoad | 4.5 | 2.097 | 3.461 | 1.647 | 282 | 25 |

SDS-PAGE analysis revealed three polypeptides of about 90, 70, and 25 kDa (*cf*. Fig. 3D) that coeluted with the enzyme activity in the fractions derived from the Superdex 200 purification step. All three bands were excised from the gel, digested with trypsin, and obtained peptides were analyzed by tandem mass spectrometry (Q-TOF). The sequences of the identified peptides were then compared with the rat reference proteome from the NCBI Protein database (Table 2). Surprisingly, the analysis indicated a lack of any proteins of unknown function. However, a thorough analysis of the available research data on the identified proteins allowed us to hypothesize that 3-hydroxybutyrate dehydrogenase type 2 (BDH2) protein might be a sought 4-oxo-L-proline reductase. BDH2 was present as a predominant protein in band C and was the only cytosolic dehydrogenase identified in analyzed protein bands. Twenty-six peptide sequences from the rat BDH2 (underlined in Fig. 2) were found in the mass spectrometry analysis to cover about 54% of its sequence. To exclude the possibility of missiHng any potential reductases due to poor extraction of tryptic peptides from the polyacrylamide gel we performed the tandem mass spectrometry identification of all proteins present in the most active fractions from the Superdex 200 purification step (F28 and F29, *cf*. Table 2). Again, BDH2 was found as the only reasonable candidate for the rat 4-oxo-L-proline reductase.

**Table 2.**
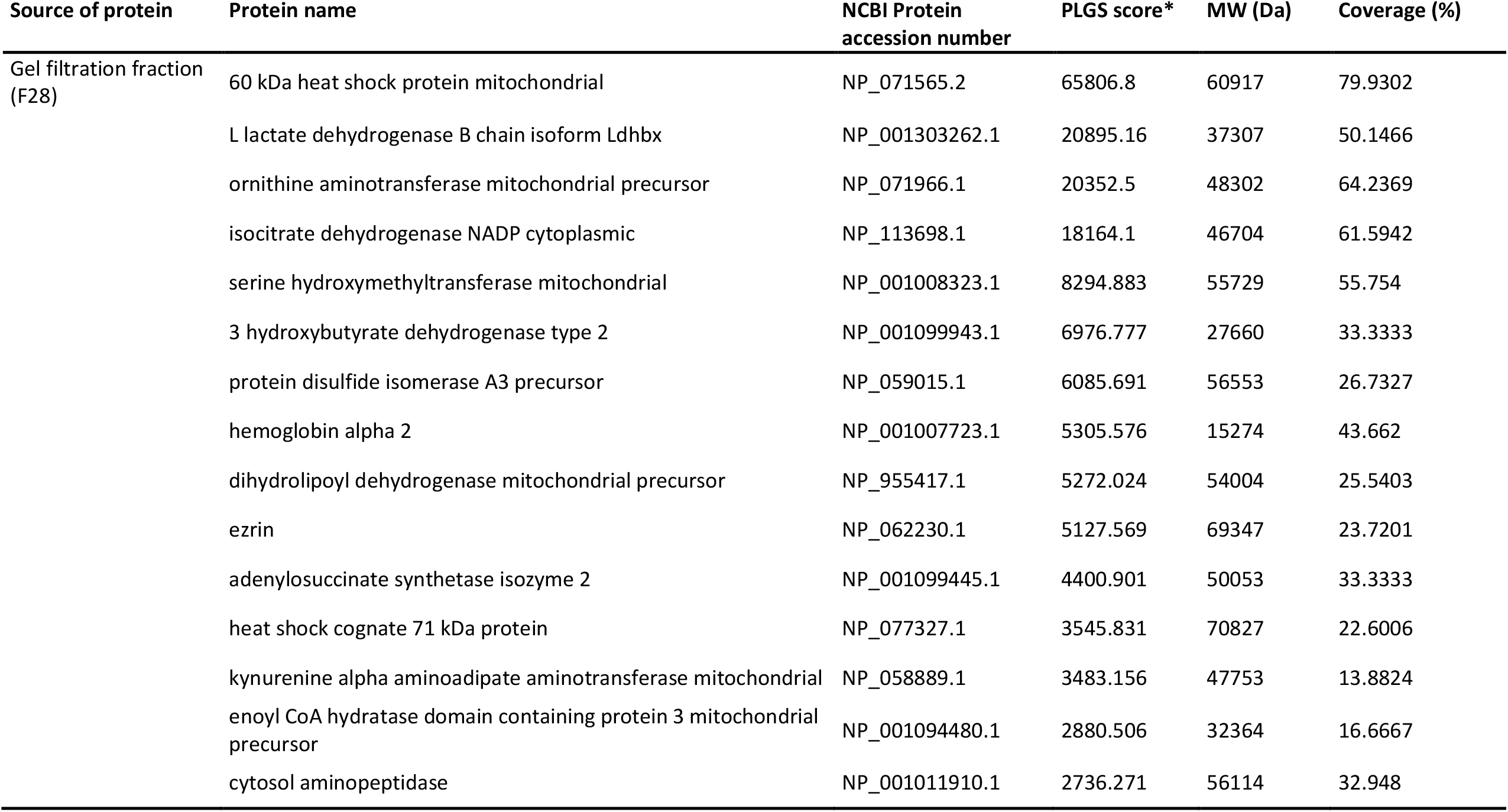
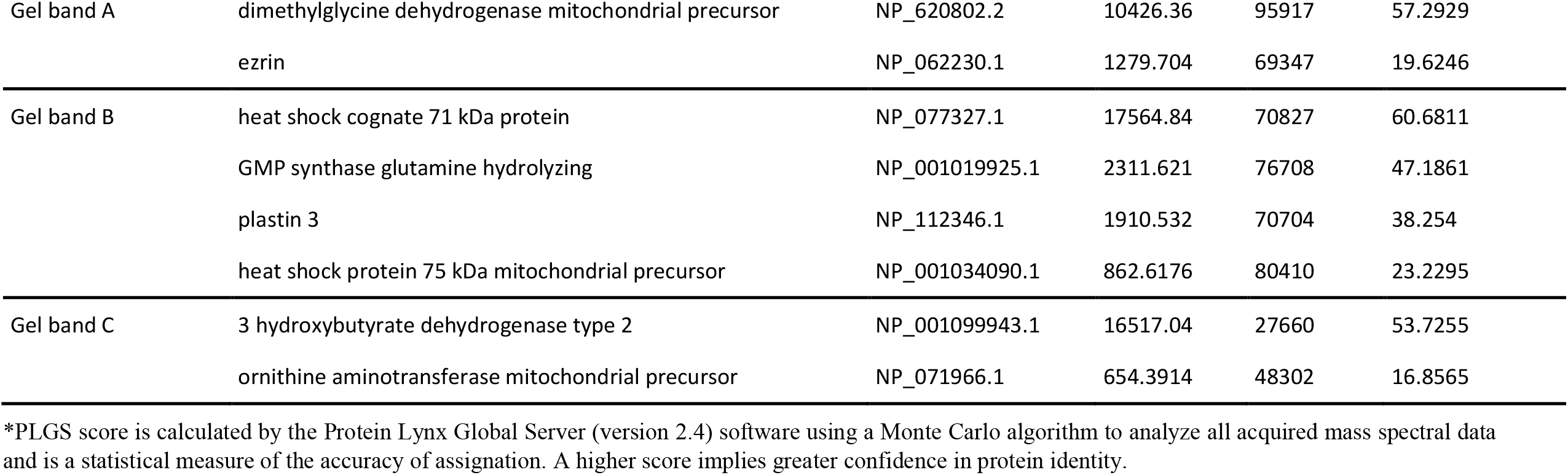
Proteins identified in fraction 28 from Superdex 200 purification step and gel bands submitted to trypsin digestion and MS/MS analysis. Identified proteins are listed according to their score as calculated using ProteinLynx Global Server software (PLGS). For each protein, the molecular weight (mW) and sequence coverage are also indicated. Occasional peptide hits corresponding to keratins have not been included.

| Source of protein | Protein name | NCBI Protein accession number | PLGS score* | MW (Da) | Coverage (%) |
| --- | --- | --- | --- | --- | --- |
| Gel filtration fraction (F28) | 60 kDa heat shock protein mitochondrial | NP_071565.2 | 65806.8 | 60917 | 79.9302 |
|  | L lactate dehydrogenase B chain isoform Ldhbx | NP_001303262.1 | 20895.16 | 37307 | 50.1466 |
|  | ornithine aminotransferase mitochondrial precursor | NP_071966.1 | 20352.5 | 48302 | 64.2369 |
|  | isocitrate dehydrogenase NADP cytoplasmic | NP_113698.1 | 18164.1 | 46704 | 61.5942 |
|  | serine hydroxymethyltransferase mitochondrial | NP_001008323.1 | 8294.883 | 55729 | 55.754 |
|  | 3 hydroxybutyrate dehydrogenase type 2 | NP_001099943.1 | 6976.777 | 27660 | 33.3333 |
|  | protein disulfide isomerase A3 precursor | NP_059015.1 | 6085.691 | 56553 | 26.7327 |
|  | hemoglobin alpha 2 | NP_001007723.1 | 5305.576 | 15274 | 43.662 |
|  | dihydrolipoyl dehydrogenase mitochondrial precursor | NP_955417.1 | 5272.024 | 54004 | 25.5403 |
|  | ezrin | NP_062230.1 | 5127.569 | 69347 | 23.7201 |
|  | adenylosuccinate synthetase isozyme 2 | NP_001099445.1 | 4400.901 | 50053 | 33.3333 |
|  | heat shock cognate 71 kDa protein | NP_077327.1 | 3545.831 | 70827 | 22.6006 |
|  | kynurenine alpha amino adipate aminotransferase mitochondrial | NP_058889.1 | 3483.156 | 47753 | 13.8824 |
|  | enoyl CoA hydratase domain containing protein 3 mitochondrial precursor | NP_001094480.1 | 2880.506 | 32364 | 16.6667 |
|  | cytosol aminopeptidase | NP_001011910.1 | 2736.271 | 56114 | 32.948 |
| Gel band A | dimethylglycine dehydrogenase mitochondrial precursor | NP_620802.2 | 10426.36 | 95917 | 57.2929 |
|  | ezrin | NP_062230.1 | 1279.704 | 69347 | 19.6246 |
| Gel band B | heat shock cognate 71 kDa protein | NP_077327.1 | 17564.84 | 70827 | 60.6811 |
|  | GMP synthase glutamine hydrolyzing | NP_001019925.1 | 2311.621 | 76708 | 47.1861 |
|  | plastin 3 | NP_112346.1 | 1910.532 | 70704 | 38.254 |
|  | heat shock protein 75 kDa mitochondrial precursor | NP_001034090.1 | 862.6176 | 80410 | 23.2295 |
| Gel band C | 3 hydroxybutyrate dehydrogenase type 2 | NP_001099943.1 | 16517.04 | 27660 | 53.7255 |
|  | ornithine aminotransferase mitochondrial precursor | NP_071966.1 | 654.3914 | 48302 | 16.8565 |

\*PLGS score is calculated by the Protein Lynx Global Server (version 2.4) software using a Monte Carlo algorithm to analyze all acquired mass spectral data and is a statistical measure of the accuracy of assignment. A higher score implies greater confidence in protein identity.

**Figure 3.**
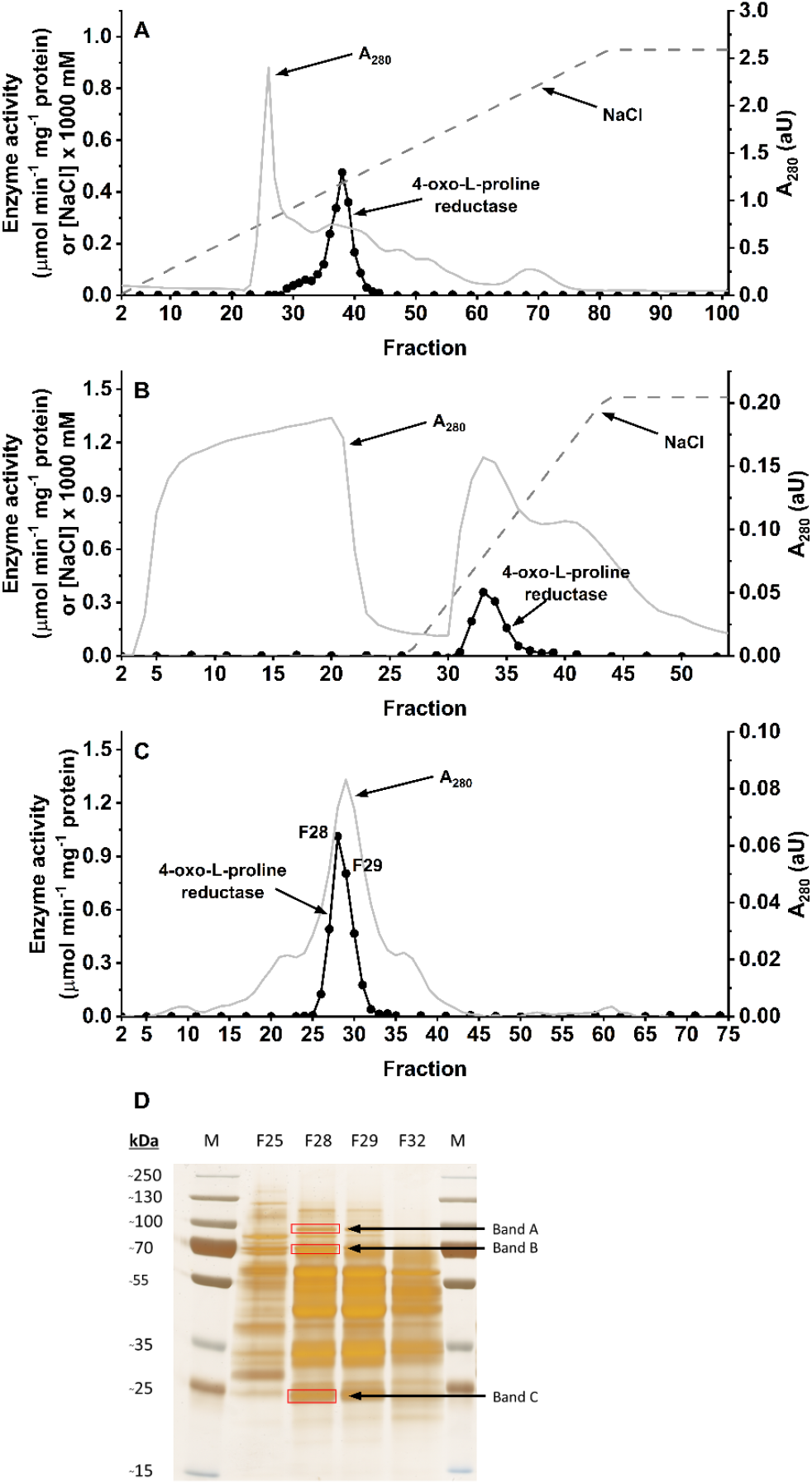
Purification of the rat 4-oxo-L-proline reductase. The enzyme was purified by column chromatography on **A**. Q Sepharose, **B**. HiScreen Blue FF, and **C**. Superdex 200 16/60 HiLoad as described in the “Experimental Procedures”section. Resulted fractions were tested for activity of 4-oxo-L-proline reductase. **D**. The indicated fractions from Superdex 200 column were analyzed by SDS-PAGE and the gel was silver-stained (24). M, prestained protein marker. Indicated bands were cut out from the gel and analyzed by mass spectrometry.

### Human and rat BDH2 catalyze the reversible reduction of 4-oxo-L-proline to *cis*-4-hydroxy-L-proline

To verify the accuracy of the molecular identification of rat BDH2 as 4-oxo-L-proline reductase as well as to compare the activity of the rat and human enzymes, both proteins were overexpressed as fusion proteins with the *N*-terminal His_6_ tag in a bacterial expression system, purified (Fig. 4) and shown to catalyze the reduction of 4-oxo-L-proline in the presence of NADH (Fig. 5). To further confirm that observed activity resulted from BDH2, and not from impurities that might co-purify with the recombinant proteins, mutated forms of both orthologues, Y157F and Y147F for rat and human, respectively, were prepared and shown to be catalytically inactive with 4-oxo-L-proline and NADH as substrates (*cf*. Fig. 2, and Fig. 5). Those results indicate that BDH2 possesses the activity of 4-oxo-L-proline reductase (EC 1.1.1.104).

**Figure 4.**
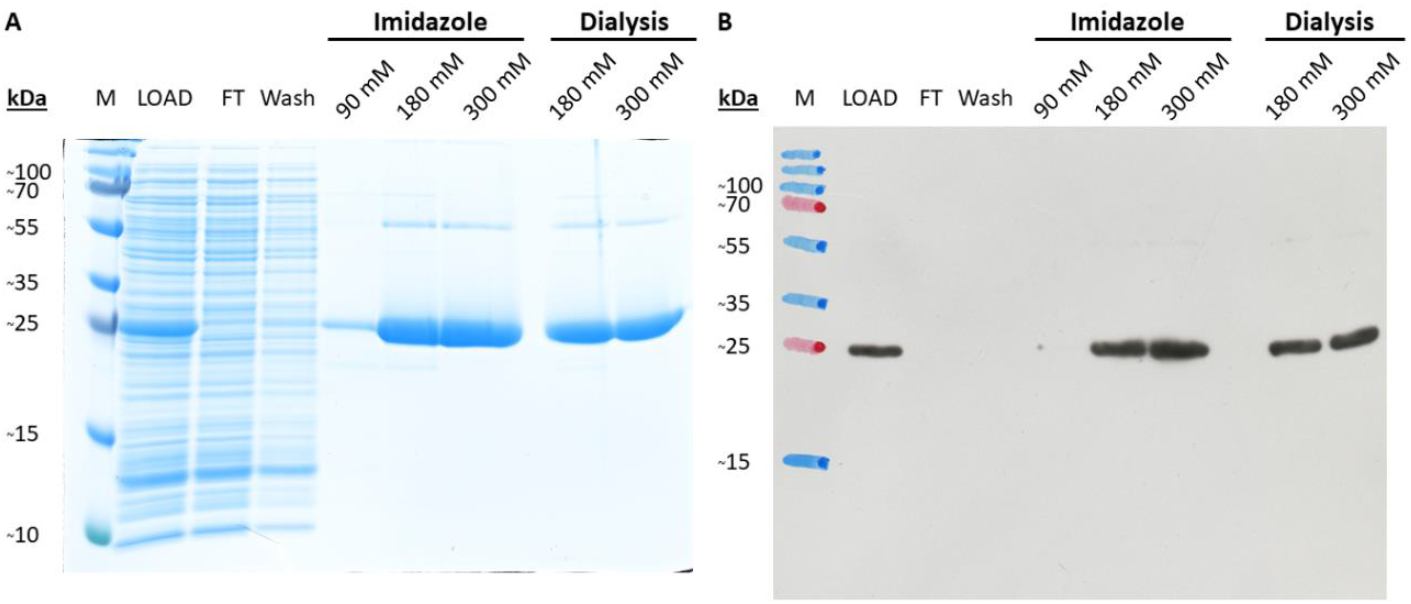
**A**. SDS-PAGE and **B**. Western blot analysis of recombinant rat BDH2 purification. Lysate of *E*.*coli* BL21 culture (LOAD) was applied to HisTrap™ FF Crude column and flow-through (FT) was collected. The column was washed with a buffer without imidazole (Wash). Retained proteins were eluted by applying the buffer with indicated concentrations of imidazole. To remove imidazole, fractions 180 mM and 300 mM were subjected to dialysis. The presence of recombinant rBDH2 protein was verified by western blot analysis using an antibody against the His_6_ tag. Analogous results were obtained for human BDH2. M, prestained protein marker copied from the blotting membrane onto the film using a felt-tip pen. The purity of both recombinant proteins was above 95%.

**Figure 5.**
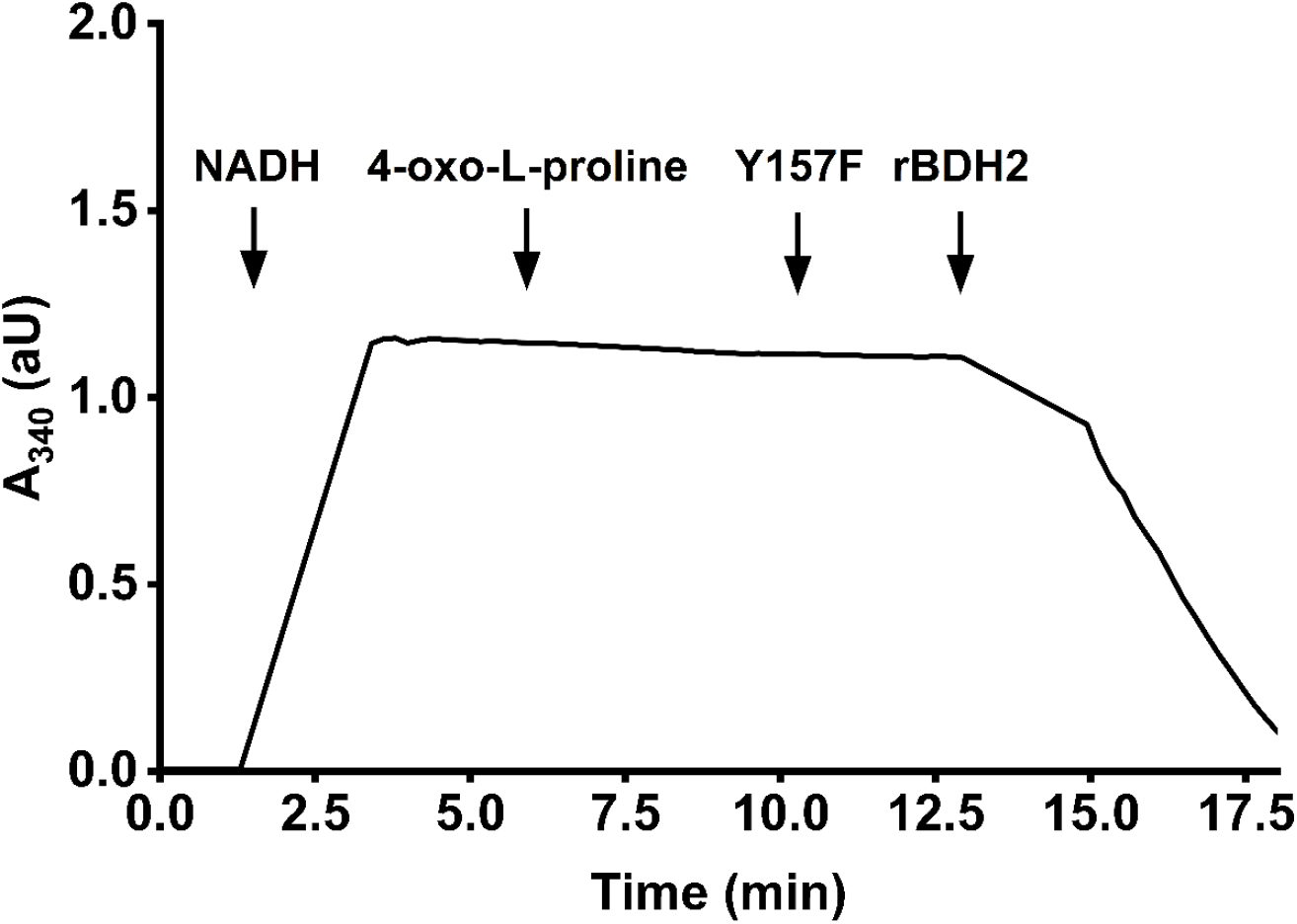
Test of the purified recombinant wild-type and mutated form of rat BDH2 activity (Y157F). The activity of the enzyme was followed spectrophotometrically by measuring the conversion of NADH into NAD^+^ (λ = 340 nm). The reaction was performed with 8 µg of Y157F protein as described in the “Experimental Procedures” section. The addition of 1.8 µg of the wild type rat BDH2 resulted in the complete oxidation of the NADH as indicated by the change in the absorbance (A_340_). An analogous result was obtained for the mutated human form of BDH2 (Y147F). The figure shows the results of a single representative assay.

Due to the high amino acid sequence and structural similarities to various bacterial hydroxybutyrate dehydrogenases, BDH2 was previously shown to be involved in the ketone body (R)-β-hydroxybutyrate metabolism (6). Furthermore, BDH2 has also been proposed to catalyze the synthesis of 2,5-dihydrobenzoic acid (2,5-DHBA, gentisic acid), putative mammalian siderophore (7). These reports led us to verify the substrate specificity of BDH2, including (R)-β-hydroxybutyrate, two isomers of 4-hydroxy-L-proline (oxidation) as well as 5-oxo-L-proline (L-pyroglutamic acid), and acetoacetate (reduction), the latter as a product of postulated (R)-β-hydroxybutyrate dehydrogenase activity of BDH2. We tested neither 3,6-dihydroxy-1,3-cyclohexadiene-1-carboxylate nor 3,6-dihydroxy-2,4-cyclohexadiene-1-carboxylate – plausible substrates for 2,5-DHBA production as these compounds have not been commercially available. Out of all tested compounds, only 4-oxo-L-proline, *cis*-4-hydroxy-L-proline, and, albeit to a much lesser extent, (R)-β-hydroxybutyrate acted as substrates for BDH2 (Table 3). Importantly, *trans*-4-hydroxy-L-proline, which has been thought to be the product of this reductase activity so far, was not oxidized to 4-oxo-L-proline in the reverse reaction, indicating a stereospecific interconversion of only *cis*-4-hydroxy-L-proline and 4-oxo-L-proline in the presence of the enzyme. Furthermore, a negligible specific activity towards (R)-β-hydroxybutyrate (0.04 ± 0.002 and 0.02 ± 0.001 µmol min^-1^ mg^-1^ protein for rBDH2 and hBDH2, respectively) comparing with the results for *cis*-4-hydroxy-(15.73 ± 0.37 and 21.16 ± 0.34 for rBDH2 and hBDH2, respectively) and 4-oxo-L-proline (22.91 ± 1.26 and 31.36 ± 0.80 for rBDH2 and hBDH2, respectively) indicate that (R)-β-hydroxybutyrate is unlikely to be a physiological substrate for BDH2.

**Table 3.** Substrate specificity of the recombinant rat and human BDH2. Activity assays were performed with the use of 2 µg of recombinant BDH2 proteins in the presence of 0.2 mM NADH or 1 mM NAD^+^ when necessary, and 2 mM concentration of the indicated substrate, with the exemption of (R)-β-hydroxybutyrate that was also tested at 50 mM concentration. Values are the means ± S.E. (error bars) of three independent experiments.

| Substrate | hBDH2 | rBDH2 |
| --- | --- | --- |
|  | *µmol min<sup>-1</sup> mg<sup>-1</sup>* | *µmol min<sup>-1</sup> mg<sup>-1</sup>* |
| 4-oxo-L-proline | 31.36 ± 0.80 | 22.91 ± 1.26 |
| 5-oxo-L-proline | N.D. | N.D. |
| acetoacetate | N.D. | N.D. |
| *trans*-4-OH-L-proline | N.D. | N.D. |
| *cis*-4-OH-L-proline | 21.16 ± 0.34 | 15.73 ± 0.37 |
| (R)-β-hydroxybutyrate | N.D. | N.D. |
| (R)-β-hydroxybutyrate (50mM)* | 0.050 ± 0.003 | 0.040 ± 0.002 |

\* 50 mM (R)-β-hydroxybutyrate was employed previously (6) as a saturating concentration of the substrate N.D. – not detectable

It was previously reported that the optimum pH value for the reaction catalyzed by 4-oxo-L-proline reductase is about 6.5 (5). To verify this information, we determined the pH range of the reductase activity (Fig. 6). Interestingly, BDH2 remained catalytically active in a broad spectrum of the pH (from 5.5 through 9.0). Moreover, the pH optimum for the rat enzyme was more evident at 6.5, while the human enzyme exhibited the highest activity in the pH range from 6.5 to 7.0, with only a slight decrease in higher pH values. These results indicate that BDH2 is most likely the same enzyme as studied by Smith and Mitoma (5).

**Figure 6.**
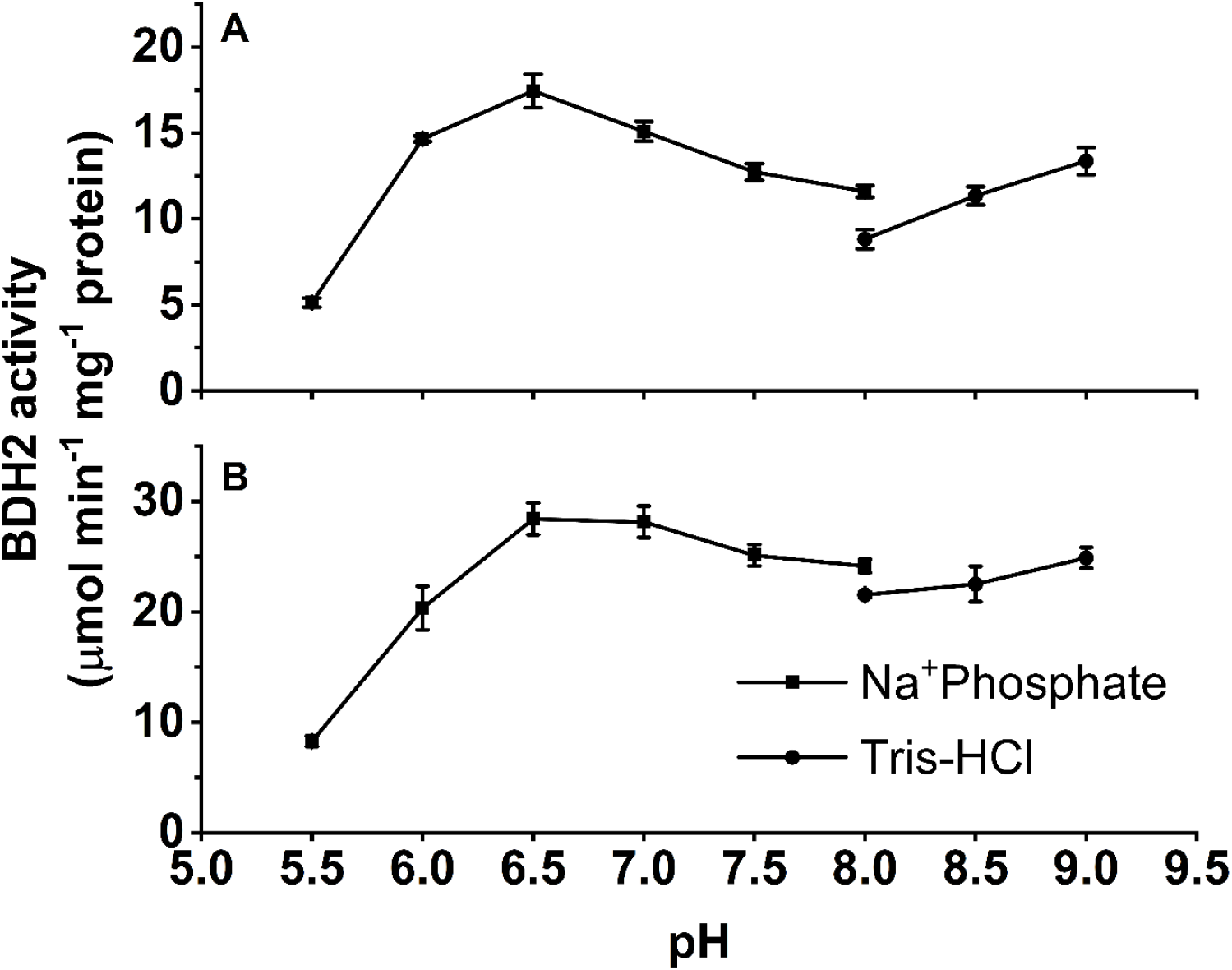
Effect of the pH on the activity of the recombinant **A**. rat and **B**. human BDH2. The reaction was performed as described in the “Experimental Procedures”.. Sodium phosphate buffer was used for the lower pH values and Tris-HCl for higher pH values. The activity of both recombinant enzymes was slightly lower in the Tris-HCl buffer compared with sodium phosphate buffer at pH = 8.0. Values are the means ± S.E. (error bars) of three independent experiments. When an error bar is not visible, the error is smaller than the width of the line.

The kinetic properties of the recombinant BDH2 proteins were investigated in detail using homogenous recombinant proteins and are compared in Table 4. Both enzymes followed the Michaelis-Menten model of enzyme kinetics. Comparing with previous studies, K_M_ values for 4-oxo-L-proline obtained in our investigation (K_M_ ≈ 0.4 – 0.5 mM) were comparable to those determined for the partially purified rabbit enzyme (K_M_ ≈ 0.6 mM) (5). In contrast, an affinity for NADH seems to be much higher than shown by Smith and Mitoma (K_M_ ≈ 840 µM) and was equal to about 2.845 ± 0.76 and 2.876 ± 0.181 µM for rat and human enzyme, respectively. These values are much more probable than the results reported previously, because of the observed saturation of the enzymes with NADH at a concentration of 50 µM.

**Table 4.**
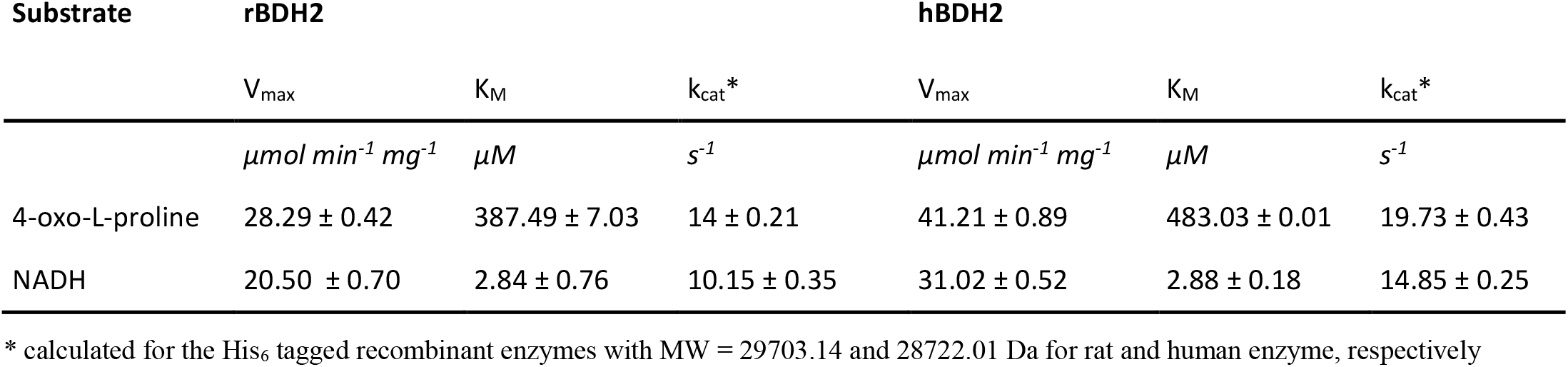
Kinetic properties of rat and human BDH2 proteins. Kinetic properties were determined with the use of purified recombinant *N*-terminal His_6_ tagged BDH2 proteins. Determinations for 4-oxo-L-proline were performed with 2 µg of the enzyme preparations in the presence of 0.2 mM NADH and variable concentrations of 4-oxo-L-proline. The measurements for NADH were performed with 0.15 µg of the enzyme preparations in the presence of 2 mM 4-oxo-L-proline and variable concentrations of NADH. Values are the means ± S.E. (error bars) of three independent experiments.

### Evidence for *cis*-4-hydroxy-L-proline as the product of BDH2 activity

So far, the reaction catalyzed by 4-oxo-L-proline reductase was only shown to produce 4-hydroxy-L-proline (4, 5), but the specific isomer of the product has never been described. To determine the stereochemistry of 4-hydroxy-L-proline generated by recombinant human BDH2, the deproteinized reaction mixture was subjected to precolumn chiral derivatization with Nα-(5-fluoro-2,4-dinitrophenyl)-l-valine amide (L-FDVA) followed by RP-HPLC and mass spectrometric analysis (8). As shown in Fig. 7, chromatographic analysis of the product revealed its perfect comigration with a commercial standard of *cis*-4-hydroxy-L-proline. The addition of *cis*-4-hydroxy-L-proline to the reaction mixture resulted in a selective increase in the peak area of the product (from 6,101 to 21,577 units) without a noticeable disturbance in its peak symmetry, confirming the product’s identity as the *cis* isomer of 4-hydroxy-L-proline. Analysis of the product by electrospray mass spectrometry indicated the presence of a protonated molecular ion with m/z 412, as expected for the L-FDVA derivative of *cis*-4-hydroxy-L-proline, which was indeed identical with that of the commercial standard of this amino acid (Fig. 7). The recorded mass spectrum of *cis*-4-hydroxy-L-proline was also in agreement with previously reported results on the L-FDVA derivative of 4-hydroxy-L-proline (8). Similar results of RP-HPLC-MS analysis were obtained for rat recombinant BDH (not shown).

**Figure 7.**
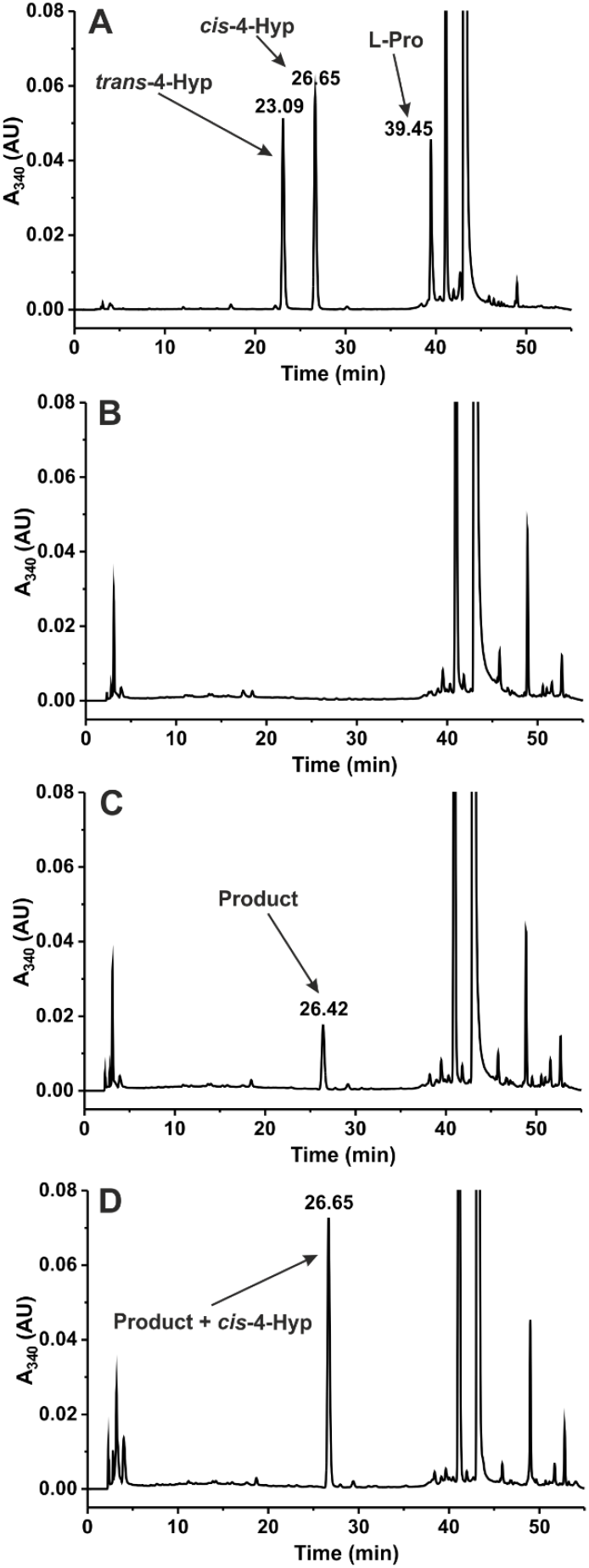
RP-HPLC analysis of the product formed by human BDH2 protein. Shown are chromatograms **A**. of standard mixture of *cis*-4-hydroxy-L-proline (*cis*-4-Hyp*), trans*-4-hydroxy-L-proline (*trans*-4-Hyp) and L-proline (0.5 nmol) after derivatization with L-FDVA; **B**. of deproteinized reaction mixtures obtained during incubation of homogenous recombinant human protein (2 µg) with 2 mM 4-oxo-L-proline and 0.2 mM NADH for 0 min or **C**. 15 min as well as **D**. following the supplementation of the former deproteinized reaction mixture with 0.5 nmol of *cis*-4-hydroxy-L-proline standard. The identity of all indicated compounds was confirmed by mass spectrometry. The sample processing and chromatographic conditions are described under “Experimental Procedures”.

These results confirm that BDH2 catalyzes the formation only of *cis*-4-hydroxy-L-proline that has been considered an exogenous and non-physiological metabolite in mammalians so far.

### HEK293T cells metabolize 4-oxo-L-proline into *cis*-4-hydroxy-L-proline

To verify whether 4-oxo-L-proline is converted into *cis*-4-hydroxy-L-proline in human cells, we investigated its metabolism in the intact HEK293T cells that express BDH2 enzyme (Fig. 9). The cells were incubated in the absence or presence of 1 mM 4-oxo-L-proline for up to 72 h. Next, samples of culture medium were withdrawn, deproteinized with perchloric acid, neutralized, and derivatized with L-FDVA. The concentrations of *trans*- and *cis*-4-hydroxy-L-proline were determined by RP-HPLC-MS, whereas the consumption of 4-oxo-L-proline was followed spectrophotometrically.

**Figure 8.**
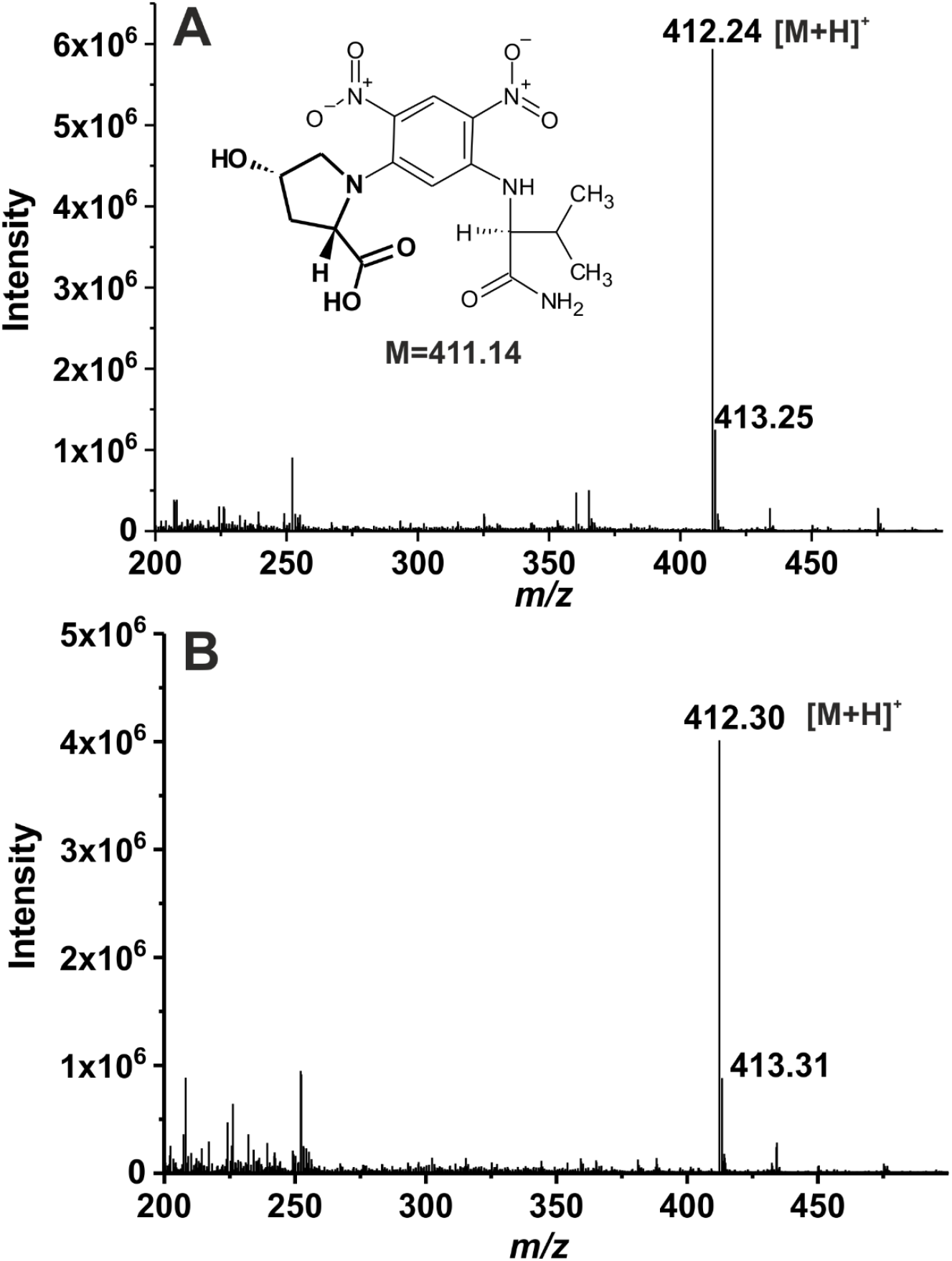
Quadrupole mass spectra of L-FDVA derivatives of *cis*-4-hydroxy-L-proline and the product formed by human BDH2 protein. The homogenous recombinant human enzyme was incubated for 15 min with 2 mM 4-oxo-L-proline and 0.2 mM NADH, and the progress of the reaction was followed spectrophotometrically at λ = 340 nm. The product was then derivatized with L-FDVA, chromatographed on a reversed-phase C18 column, and analyzed by mass spectrometry. Mass spectra, covering the mass range *m/z* 200–500, **A**. of commercial *cis*-4-hydroxy-L-proline derivatized with L-FDVA and **B**. the product generated by human BDH2 enzyme were acquired. The structure of the L-FDVA derivative of *cis*-4-hydroxy-L-proline (in bold) is also shown.

**Figure 9.**
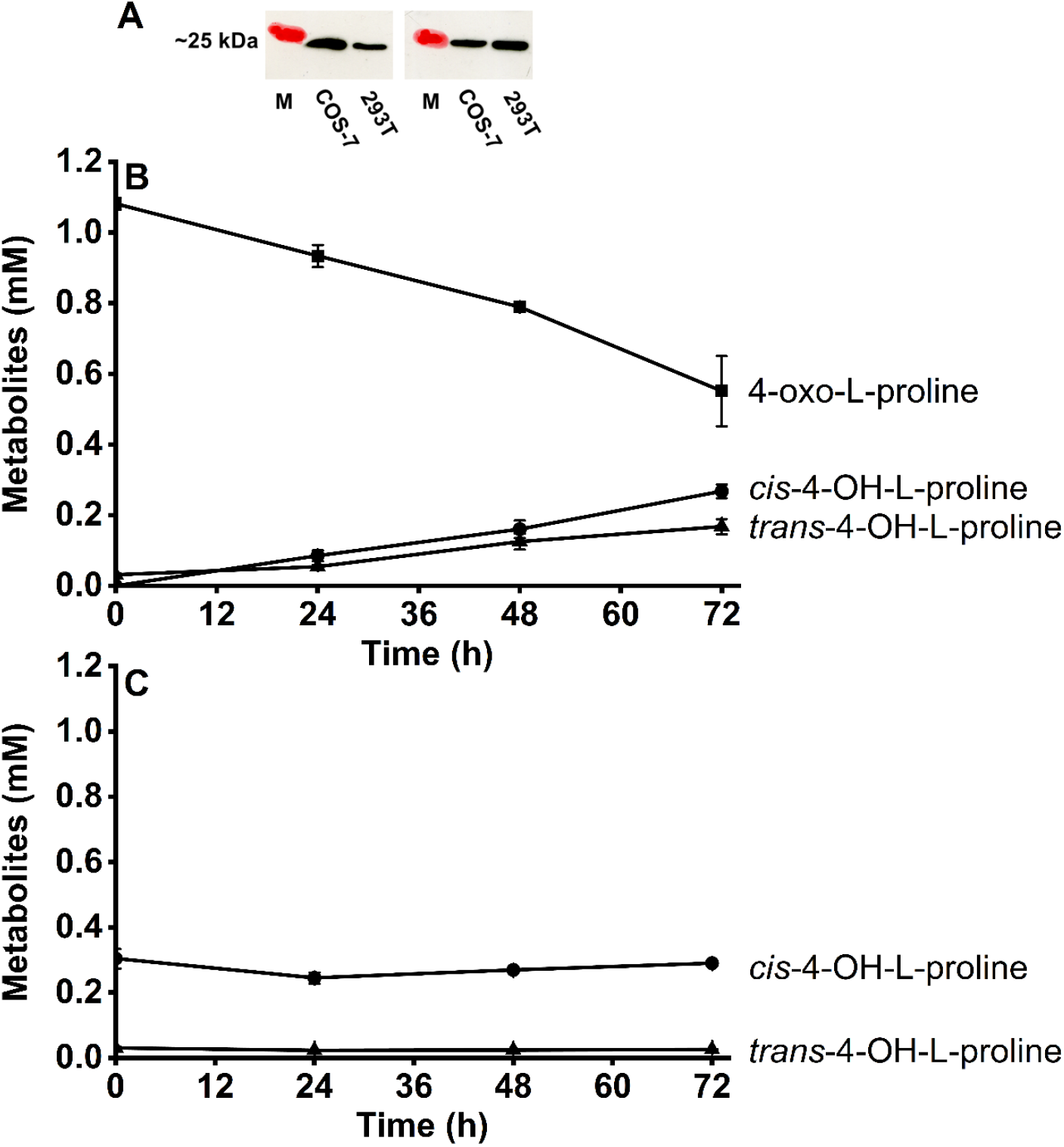
Metabolism of 4-oxo-L-proline and *cis*-4-hydroxy-L-proline in HEK293T cells. **A**. Western blot analysis of HEK293T and COS7 cell extracts from two different passages, showing the endogenous expression of BDH2. The analysis was performed using 30 µg of total protein with a polyclonal rabbit antibody against human BDH2. M, prestained protein marker copied from the blotting membrane onto the film using a felt-tip pen; **B**. Changes in the extracellular concentration of 4-oxo-L-proline and its metabolites in HEK293T cells, following supplementation of the cell culture medium with 1 mM 4-oxo-L-proline. **C**. Changes in the extracellular concentration of *trans*- and *cis*-4-hydroxy-L-proline in HEK293T cells, following supplementation of the cell culture medium with 0.3 mM *cis*-4-hydroxy-L-proline. The cells were plated in 6-well dishes and grown for 24 h. After that time, the cell culture medium was supplemented with the indicated amino acids, and the incubation was continued for up to 72 h as described under “Experimental Procedures”. Values are the means ± S.E. (error bars) of three independent experiments performed with cells from three different culture passages (n = 3). When no error bar is shown, the error is smaller than the width of the line.

As shown in Fig. 9, HEK293T constantly took up 4-oxo-L-proline from the culture medium and converted it into *cis*-4-hydroxy-L-proline that was then released back into the extracellular milieu. No formation of *cis*-4-hydroxy-L-proline was detected in the absence of 4-oxo-L-proline (not shown). After 72h of incubation, the extracellular concentration of 4-oxo-L-proline dropped by ≈ 0.5 mM, whereas the *cis*-4-hydroxy-L-proline accumulated up to ≈ 0.3 mM. More intriguingly, the formation of *cis* isomer was accompanied by an accumulation of the *trans* one at up to ≈ 0.2 mM concentration. These results suggested that a considerable fraction of *cis*-4-hydroxy-L-proline might have been converted into the *trans* isomer, a physiological metabolite in mammalians. Alternatively, 4-oxo-L-proline might have exerted an inhibitory action on the breakdown of the endogenous *trans*-4-hydroxy-L-proline. The substoichiometric production of *cis*-4-hydroxy-L-proline would then reflect its retention in the intracellular spaces accompanied by its incorporation into cell proteins (9). To verify these hypotheses, we investigated the metabolic fate of the *cis* isomer in HEK293T. As shown in Fig. 9, the cells did neither convert *cis*-4-hydroxy-L-proline into *trans*-4-hydroxy-L-proline nor accumulate the latter metabolite in the culture medium, indicating both a lack of *cis*-*trans* isomerization of 4-hydroxy-L-proline and any impact of the *cis* isomer on the metabolism of the endogenous *trans*-4-hydroxy-L-proline. These findings imply that 4-oxo-L-proline acts as an inhibitor of the *trans*-4-hydroxy-L-proline breakdown in HEK293T cells.

A routine daily inspection of HEK293T cultures indicated that the cells incubated in the presence of 4-oxo-L-proline might show a higher death rate than the control ones. To test this possibility, the viability of both control and 4-oxo-L-proline-treated cells was determined using the MTT assay.

As shown in Fig. 10, prolonged incubation of the cells with the amino acid led to a decrease in their viability by ≈ 50% in comparison with that determined for the control cells, suggesting an antiproliferative and cytotoxic effect of *cis*-4-hydroxy-L-proline as reported previously (10, 11). A similar impact on the cell viability was also found with African green monkey COS-7 cells (not shown).

**Figure 10.**
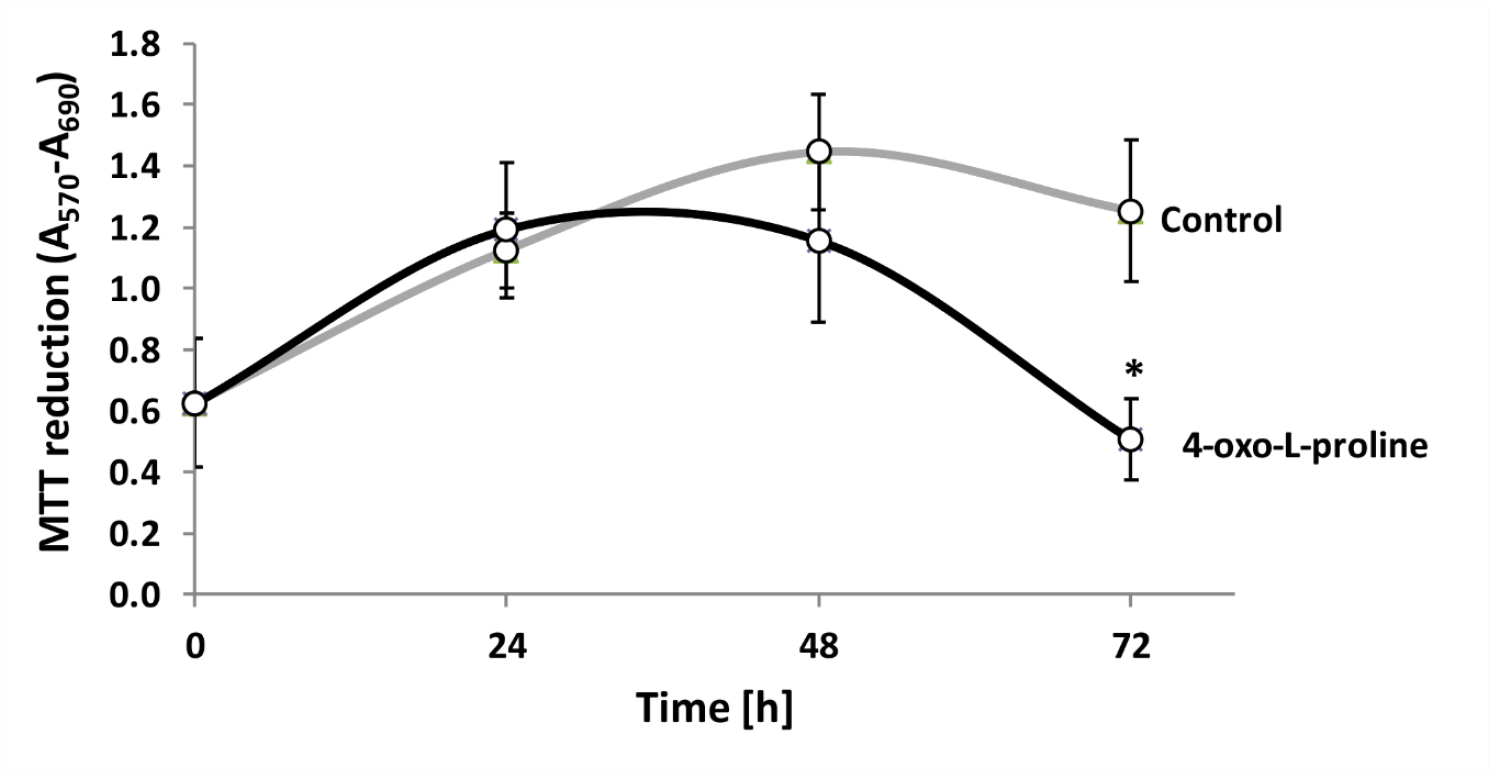
The cell viability of HEK293T cells in the presence of 4-oxo-L-proline. Cell viability was tested at the indicated time points by the MTT assay. The cells were seeded in 12-well dishes and grown for 24 h. After that time, the cell culture medium was replaced with the fresh one supplemented with 1 mM 4-oxo-L-proline, and the incubation was continued for up to 72 h as described under “Experimental Procedures”. Values are the means ± S.E. (error bars) of three independent experiments performed in triplicates with cells from three different culture passages (n = 3). The data were assumed to be distributed normally. Statistical significance was analyzed using a one-tailed paired Student’s *t*-test. ^*^p < 0.02.

Taken together, these results indicate that BDH2 enzyme is operational in the intact HEK293T cells, yielding *cis*-4-hydroxy-L-proline, a non-physiological and potentially toxic metabolite, whereas 4-oxo-L-proline is most likely an inhibitor of *trans*-4-hydroxy-L-proline metabolism in human cells.

## Discussion

Although the reaction catalyzed by 4-oxo-L-proline reductase is currently considered a normal part of metabolic pathways for L-proline degradation in mammalians, as depicted in the Kyoto Encyclopedia of Genes and Genomes (KEGG) for example (12), the identity of this enzyme and its biological impotence are still unknown. Here, we report the identification of rat 4-oxo-L-proline reductase as BDH2, disclosing the identity of the mammalian enzyme. This conclusion is based on the following findings: (i) a multistep purification of the 4-oxo-L-proline-reducing activity from rat kidneys resulted in the identification of the protein BDH2 as the only meaningful candidate for the enzyme; (ii) the recombinant rat and human BDH2 catalyze the NADH-dependent reduction of 4-oxo-L-proline, yielding *cis*-4-hydroxy-L-proline, whereas mutants of these enzymes harboring mutations of the key catalytic residues (Y157F and Y147F, respectively) are catalytically inactive; and (iii) the identity of the product made by the recombinant enzymes was confirmed by both reversed-phase chromatography and mass spectrometry. The identification of 4-oxo-L-proline reductase as BDH2 is also consistent with the findings that HEK293T cells endogenously expressing BDH2 protein efficaciously convert 4-oxo-L-proline to *cis*-4-hydroxy-L-proline, whereas no other enzyme has been shown to generate the *cis* isomer of 4-hydroxy-L-proline in vertebrates yet (for review see Ref. 13).

BDH2 has a typical NAD(H) binding Rossmann-fold domain in its structure and belongs to the short-chain dehydrogenase/reductase (SDR) superfamily. In humans, enzymes of this cluster are involved in the metabolism of a large variety of compounds, including steroid hormones, lipids, and xenobiotics (14). BDH2 was initially reported as a putative cytosolic β-hydroxybutyrate dehydrogenase contributing to the oxidation of the ketone body β-hydroxybutyrate (6). But the enzyme was shown to be very poor in oxidizing β-hydroxybutyrate, with a K_M_ of 12 mM and k_cat_ value of 1 min^-1^ at 50 mM concentration of the substrate, which also finds confirmation in the present study. Such a low turnover number of the enzyme, resembling that for an enzyme of posttranslational modification rather than for a metabolic one (15), clearly pleads against β-hydroxybutyrate being its physiological substrate. Indeed, no disturbances in ketone bodies metabolism were found in BDH2 deficient mice (16). Alternatively, BDH2 was proposed to be the homolog of bacterial EntA protein, which reduces 2,3-dihydro-2,3-dihydroxybenzoic acid (2,3-diDHBA) to 2,3-dihydroxybenzoic acid (2,3-DHBA), and to catalyze the synthesis of 2,5-dihydroxybenzoic acid (2,5-DHBA, gentisic acid), a putative mammalian siderophore (7). Intriguingly, BDH2 has never been shown in a direct experiment (*i*.*e. in vitro*) to catalase the production of 2,5-DHBA and instead of this, the formation of 2,3-DHBA from 2,3-diDHBA in the presence of the enzyme was evidenced (7). It is also unclear which metabolic pathway could provide a putative substrate for the BDH2-dependent formation of 2,5-DHBA in mammalians. For these and other factual reasons, the physiological role of 2,5-DHBA and the importance of BDH2 for its synthesis were questioned experimentally by others (17).

We show in the present work that BDH2 catalyzes the reversible reduction of 4-oxo-L-proline to *cis*-4-hydroxy-L-proline, with a high substrate specificity and catalytic activity. The formation of *cis*-4-hydroxy-L-proline is remarkable as, to our knowledge, no enzyme has been shown to catalase the production of the *cis* isomer in vertebrate species to date. This finding also reveals that the irreversibility of this reaction reported by Smith and Mitoma (5) was only apparent, and resulted from the use of *trans*-4-hydroxy-L-proline in enzymatic tests. Concerning the plausible substrates, neither 5-oxo-L-proline nor *trans*-4-hydroxy-L-proline was accepted, indicating a preferential requirement for the presence of an OH/keto group at carbon 4 position and specific spatial orientation of 4 OH group in a reduced substrate. The catalytic efficiency (k_cat_/K_M_) of the human enzyme on 4-oxo-L-proline (2450 min^-1^ × mM^-1^) was 9400-fold higher than that on (R)-β-hydroxybutyrate (0.26 min^-1^ × mM^-1^, 6), confirming that the latter compound is unlikely to be a physiological substrate. Whereas a very weak activity towards (R)-β-hydroxybutyrate and no activity on acetoacetate imply that cyclic compounds are much better substrates for BDH2 than the linear ones. Unfortunately, no information on the specific activity and the kinetic parameters of BDH2 for 2,3-diDHBA was reported by Devireddy and colleagues (7), hence it is impossible to compare the catalytic activity of the enzyme on 2,3-diDHBA to that determined for 4-oxo-L-proline.

The results presented here identified 4-oxo-L-proline as a novel substrate for BDH2. Although this amino acid is a poorly studied metabolite, it is occasionally detected in metabolomics experiments as a constituent of both human plasma (3, 18) and cells (2, 19). The origin of 4-oxo-L-proline is currently unclear, but the finding that its intracellular concentration doubled in human T cells incubated with a high concentration of L-arginine suggests that 4-oxo-L-proline is indeed an endogenous metabolite, most likely contributing to L-arginine/L-proline metabolic pathways (19). Also, 4-oxo-L-proline might come from food as it is detected in dairy products for example (20). An alternative explanation would be that 4-oxo-L-proline is endogenously produced from *cis*-4-hydroxy-L-proline in the reverse reaction catalyzed by BDH2. This enzyme is certainly capable to catalyze such a reaction *in vitro* in the presence of NAD^+^ at a 1 mM concentration that is well within the physiological values reported in human cells (0.2 – 1 mM, 21). But in contrast to this reflection, no conversion of *cis*-4-hydroxy-L-proline into 4-oxo-L-proline was detected in HEK293T cells incubated with the former compound, suggesting that the reduction of 4-oxo-L-proline is the preferred reaction *in vivo*.

The incubation of HEK293T cells in the presence of 4-oxo-L-proline resulted not only in its conversion to *cis*-4-hydroxy-L-proline but also in a considerable accumulation of *trans*-4-hydroxy-L-proline in the culture medium; whereas no production of *trans*-4-hydroxy-L-proline was detected in cells incubated with *cis*-4-hydroxy-L-proline alone. This indicates that 4-oxo-L-proline is most likely an inhibitor of hydroxyproline dehydrogenase (Proline dehydrogenase 2, PRODH2) that converts *trans*-4-hydroxy-L-proline to delta-1-pyrroline-3-hydroxy-5-carboxylate, hence initiating the degradation of the endogenous *trans*-4-hydroxy-L-proline. Thus, 4-oxo-L-proline apparently blocks the degradation pathway of *trans*-4-hydroxy-L-proline in HEK293T cells, resulting in its accumulation in the culture medium. This effect in fact resembles the biochemical phenotype associated with the deficiency of hydroxyproline dehydrogenase, a benign metabolic disorder (22). Interestingly, several analogs of *trans*-4-hydroxy-L-proline were tested as potential inhibitors of PRODH2, including *trans*-4-hydroxy-L-proline, *cis*-4-hydroxy-L-proline, and *cis*-4-hydroxy-D-proline, but none of them caused inhibition at a concentration as high as 5 mM (23). Further studies are thus needed to evaluate the effect of 4-oxo-L-proline and its analogues on PRODH2 activity as inhibitors of this enzyme could have therapeutic value in the primary hyperoxalurias (23).

In conclusion, we have shown here that mammalian BDH2 is 4-oxo-L-proline reductase, an enzyme catalyzing reversible reduction of 4-oxo-L-proline to *cis*-4-hydroxy-L-proline, and not to *trans*-4-hydroxy-L-proline as currently believed. BDH2 is therefore the first enzyme capable of producing the *cis* isomer of 4-hydroxy-L-proline to be identified in vertebrates. BDH2 also allows intact HEK293T cells to metabolize 4-oxo-L-proline to *cis*-4-hydroxy-L-proline which is now thought an exogenous metabolite in vertebrates. Finally, this work also shows that 4-oxo-L-proline may act as an inhibitor of the *trans*-4-hydroxy-L-proline breakdown in human cells.

## Experimental procedures

### Materials

Reagents, of analytical grade whenever possible, were from Sigma or Merck (Darmstadt, Germany). 4-oxo-L-proline hydrobromide was purchased from Alfa Aesar (90+% purity, Ward Hill, USA) or Fluorochem (95% purity, Hadfield, UK). Q-Sepharose FF resin, HiScreen Blue FF, Superdex 200 16/60 HiLoad, and HisTrap FF crude columns were obtained from GE Healthcare Bio-Sciences (Uppsala, Sweden). Vivaspin 20 centrifugal concentrators were from Sartorius (Stockport, UK). Enzymes and DNA modifying enzymes were obtained from Thermo-Fermentas (Waltham, USA), A&A Biotechnology (Gdynia, Poland), or Bio-Shop (Burlington, Canada).

### Assay of 4-oxo-L-proline reductase activity

The enzyme activity was determined by the modified method employed previously (5). Briefly, the activity was followed spectrophotometrically at 37°C by measuring the rate of NADH conversion into NAD^+^, which is accompanied by a decrease in absorbance at λ = 340 nm (ε = 6.22 mM^-1^ cm^-1^). The standard incubation mixture (1 ml) contained 80 mM Na^+^Phosphate, pH 6.5; 1 mM DTT; 0.2 mM NADH; 2 mM 4-oxo-L-proline. The latter one was prepared as a fresh 25 mM solution and pH adjusted to 6.0 with 1M NaHCO_3_ and filtered using 0.22 µm Spin-X cellulose acetate centrifuge tube filters (Costar, USA). The actual concentration of 4-oxo-L-proline in the stock solution was verified spectrophotometrically. The reaction was started by the addition of the enzyme preparation and carried out at 37°C for 15 min unless otherwise described. Kinetic properties and substrate specificity of the BDH2 enzymes were determined in the standard incubation mixture supplemented with 0.1 mg/ml bovine serum albumin. Oxidase activity of both enzymes was determined in the analogical buffer with 50 mM Tris-HCl, pH 9.0 in the presence of 1 mM NAD^+^ instead of 0.2 mM NADH. All reactions were linear for at least 10 min under all studied conditions.

### Purification of rat 4-oxo-L-proline reductase

Eighteen male WAG rats, aged 3 months, were purchased from the Animal House of the Mossakowski Medical Research Centre, Polish Academy of Sciences (Warsaw, Poland). The animals were euthanized by a carbon dioxide euthanasia (Directive 2010/63/EU of the European Parliament). Rat kidneys (40 g) were homogenized in a Waring Blender 7011HS (4 cycles × 30 sec with 10-sec pause) with three volumes (w/v) of a buffer consisting of 20 mM Tris-HCl, pH 8.0, 1 mM DTT, 20 mM KCl, 4 µg/ml leupeptin. The homogenate was centrifuged for 20 min at 15,000 × g at 4°C. The resulting supernatant (120 ml) was then fractionated between 0% and 10% concentration (w/v) of polyethylene glycol 4000. After 15 min incubation on ice, the sample was centrifuged for 10 min at 15,000 × g at 4°C. The supernatant was again submitted for fractionation with PEG-4000 concentration (w/v) between 10% and 25%. After 15 min incubation on ice, the sample was centrifuged for 10 min at 15,000 × g at 4°C. The 10-25% precipitate was dissolved in 120 ml of homogenization buffer and frozen at -70°C before purification.

Clarified sample 10-25% was applied to a Q Sepharose column (100 ml) equilibrated with buffer A consisting of 20 mM Tris-HCl, pH 8.0, 1 mM DTT, 20 mM KCl, 1 µg/ml leupeptin. The column was washed with 240 ml of buffer A, developed with a linear NaCl gradient (0-1 M in 400 ml) in buffer A, and fractions (5 ml) were collected. The most active fractions from the Q Sepharose column were pooled (20 ml), diluted to 54 ml with buffer C (80 mM Na^+^Phosphate, pH 6.5, 1 mM DTT, 20 mM KCl, 0.5 mM phenylmethylsulfonyl fluoride (PMSF), and applied to a HiScreen Blue FF column (4.7 ml) equilibrated with buffer C. The column was washed with 56 ml of buffer C, developed with a linear NaCl gradient (0-1.5 M in 51 ml) in buffer C, and fractions (3 ml) were collected. The most active fractions from the HiScreen Blue FF column (12 ml) were pooled, concentrated to 4.5 ml using Vivaspin 20 ultrafiltration unit, and 2 ml of the sample was loaded on a Superdex 200 16/60 HiLoad column (120 ml) equilibrated with buffer D consisting of 80 mM Na^+^Phosphate, pH 6.5, 1 mM DTT, 20 mM KCl, 0.1 M NaCl. The gel filtration column was then developed with 140 ml of buffer C and 2 ml fractions were collected. All purification steps were performed at 4°C and the enzymatic preparation was stored at -70°C between steps.

### Identification of the rat 4-oxo-L-proline reductase by tandem mass spectrometry

The protein bands of the most active fractions co-eluting with 4-oxo-L-proline reductase activity in the Superdex 200 16/60 HiLoad purification step were cut from a 10% polyacrylamide SDS gel and digested with trypsin. Appropriate negative controls from two fractions lacking the activity were prepared as well. In-gel digestions of the peptides were performed as described previously (24). Peptides were analyzed by nanoUPLC-tandem mass spectrometry employing Acquity nanoUPLC coupled with a Synapt G2 HDMS Q-TOF mass spectrometer (Waters, Milford, USA) fitted with a nanospray source and working in MS^E mode under default parameters. Briefly, the products of in-gel protein digestion were loaded onto a Waters Symmetry C18 trapping column (20 mm × 180 µm) coupled to the Waters BEH130 C18 UPLC column (250 mm x 75 µm). The peptides were eluted from these columns in a 1-85% gradient of acetonitrile in water (both containing 0.1% formic acid) at a flow rate of 0.3 µl/min. The peptides were directly eluted into the mass spectrometer. Data were acquired and analyzed using MassLynx 4.1 software (Waters, USA) and ProteinLynx Global Server 2.4 software (PLGS, Waters, USA) with a False Discovery Rate ≤ 4%. To identify 4-oxo-L-proline reductase, the complete rat (*Rattus norvegicus*) reference proteome was downloaded from the NCBI Protein database, randomized, and used as a databank for the MS/MS software.

### Overexpression and purification of the recombinant BDH2 proteins and inactive mutants

Rat total RNA was prepared from 100 mg of kidneys with the use of TriPure reagent (Roche, Switzerland) according to the manufacturer’s instructions. cDNA was synthesized using Moloney murine leukemia virus reverse transcriptase (TranScriba, A&A Biotechnology, Poland), with oligo(dT)_18_ primer and 2.5 mg total RNA according to the manufacturer’s instructions. The open reading frame (ORF) encoding human enzyme was purchased from DNASU Plasmid Repository (cloneID: HsCD00640151). The open reading frames encoding rat (NCBI Reference Sequence: NM_001106473.1) and human (NM_020139.4) BDH2 protein were PCR-amplified using Pfu DNA polymerase. Both rat and human ORFs coding for BDH2 were amplified using specific 5’ primers containing the initiator codon included in the NdeI site and 3’ primers containing the stop codon flanked by a KpnI site (for primer sequence refer to table 5).

**Table 5.**
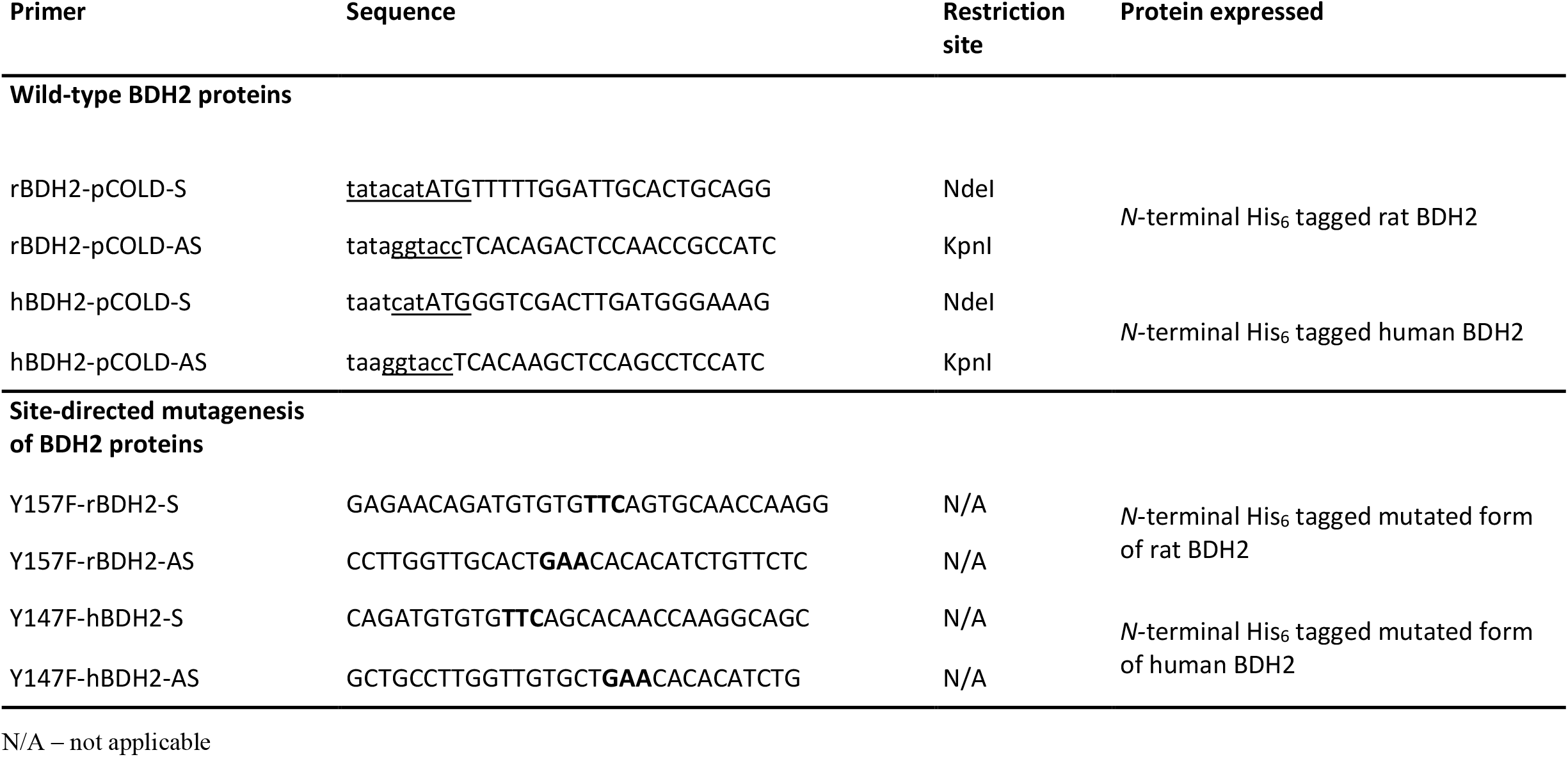
Sequences of primers used for PCR amplification of BDH2 ORFs and site-directed mutagenesis experiments. The nucleotides corresponding to the coding sequences are in capital letters, restriction sites are underlined, and mutated codons are shown in boldface type.

The amplified DNA products of the expected size were digested with the appropriate restriction enzymes and cloned into the pCOLD I expression vector (Takara Bio, Kusatsu, Japan), which allows for the production of proteins with an *N*-terminal His_6_ tag. All constructs were verified by DNA sequencing.

For protein production, *E. coli* BL21(DE3) cells were transformed with an appropriate DNA construct and a single colony was selected to start an over-night pre-culture. 300 mL of LB broth (with 100 mg/mL ampicillin) was inoculated with 30 ml of the pre-culture and incubated at 37°C and 175 rpm until an OD_600_ of 0.5 was reached. The culture was placed on ice for 20 min (cold-shock) to induce protein expression. Cells were incubated for 16 h at 13°C, 175 rpm, and harvested by centrifugation (6000 × g for 10 min). The cell paste was suspended in 15 ml lysis buffer consisting of 80 mM Na^+^Phosphate, pH 6.5, 1 mM DTT, 20 mM KCl, 1 mM PMSF, 5 µg/ml leupeptin, 5 µg/ml antipain, 0.25 mg/ml hen egg-white lysozyme (BioShop, Canada) and 250 U of Viscolase (A&A Biotechnology, Poland). The cells were lysed by freezing in liquid nitrogen and, after thawing and vortexing, the extracts were centrifuged at 4°C (20,000 × g for 20 min).

For the purification of recombinant BDH2 proteins, the supernatant of *E. coli* lysate (15 ml) was diluted 3-fold with buffer A (100 mM Na^+^Hepes pH 8.0, 200 mM NaCl, 30 mM imidazole, 1 µg/ml leupeptin, 1 µg/ml antipain) and applied onto a HisTrap FF crude column (1 ml) equilibrated with the same buffer. The column was then washed with 10 ml buffer A, and the retained proteins were eluted with a stepwise gradient of imidazole (5 ml of 90 mM, 5 ml of 180 mM, and 5 ml of 300 mM) in buffer A. The recombinant proteins were present in both 180 mM and 300 mM imidazole fractions. Both fractions exhibited > 95% purity as confirmed by SDS-PAGE. The enzyme preparation was submitted to dialysis against buffer consisted of 80 mM Na^+^Phosphate, pH 7.5, 1 mM DTT, 100 mM NaCl, 6% sucrose, 1 µg/ml leupeptin, 1 µg/ml antipain. The purified proteins were aliquoted and stored at -70°C.

Mutated forms of rat and human BDH2 enzymes (Y157F and Y147F, respectively) were generated by site-directed mutagenesis using a QuikChange II XL kit (Stratagene, La Jolla, CA, USA), with pCOLD I/rBDH2 and pCOLD I/hBDH2 plasmids as templates and mutagenic primers Y157F-rBDH2-S and Y157F-rBDH2-AS as well as Y147F-hBDH2-S and Y147F-hBDH2-AS, respectively (see table 5). Both mutated forms of BDH2 were then produced in *E. coli* BL21(DE3) and purified using HisTrap FF crude column (1 ml) as described for the wild type enzymes (data not shown).

### Product analysis

#### Products of enzymatic reaction

To obtain products formed in the reactions catalyzed by recombinant BDH2 proteins for mass spectrometry analysis, 2 µg of the homogenous recombinant enzyme was incubated in the reaction mixture (1 ml) containing 10 mM Na^+^Phosphate, pH 6.5; 1 mM DTT; 0.2 mM NADH, and 2 mM 4-oxo-L-proline. The reaction was started by the addition of BDH2 and followed up spectrophotometrically as described above. After 0 and 15 min at 37°C, 0.9 ml of the reaction mixture was removed and mixed with 200 µl of ice-cold 10% (w/v) HClO_4_ to stop the reaction. Samples were centrifuged at 13,000 × g for 10 min at 4°C and the supernatants (1.1 ml) were immediately withdrawn and neutralized with 70 µl of 3 M KOH/3M KHCO_3_. The salt precipitate was removed by centrifugation (13,000 × g for 10 min). The clear supernatants were subjected to precolumn chiral derivatization of 4-hydroxy-L-proline isomers with Nα-(5-fluoro-2,4-dinitrophenyl)-l-valine amide (L-FDVA), whereas the separation of derivatized amino acids was accomplished by RP-HPLC according to a slightly modified method described by Langrock and coworkers (8). Briefly, twenty-five microliters of neutralized supernatant were mixed with 10 µl of 1M NaHCO_3_, followed by 40 µl of 35 mM L-FDVA (dissolved in acetone). The mixture was incubated for 90 min at 40°C, and the reaction was stopped by the addition of 10 µl 1M HCl. Next, the mixture was diluted with 50 µl acetonitrile and 115 µl water, and the samples were stored at -20°C before further analysis. The amino acid derivatives were separated in a gradient mode on the Zorbax SB-C18 column (ODS, 4.6 × 250 mm, 5-µm particle size) using Waters HPLC 600 system equipped with Waters 2487 UV detector and Waters ZQ quadrupole mass spectrometer fitted with an electrospray source. Mobile phases consisted of solvent A, containing 0.1% formic acid in the water, and solvent B, containing 0.1% formic acid in acetonitrile. The separation was performed in a linear gradient from 20 to 30% of solvent B for 30 min and subsequently, from 30 to 65% for 19 min at a flow rate of 1 ml/min, followed by the column equilibration for a further 11 min under the initial conditions. The column eluate was monitored by the UV detector at λ = 340 nm, followed by the mass spectrometer, operating in positive electrospray ionization-MS mode. The mass spectral data were recorded for *m/z* = 200-500 to detect L-FDVA and derivatives of L-proline and 4-hydroxy-prolines. The ESI-MS source was set at a temperature of 90°C, the capillary voltage of 3.0 kV, and cone voltage of 40 V. The flow rate of the desolvation gas (nitrogen) was 900 liters/h. Quantification was achieved using external standards of all studied amino acids, after their derivatization with L-FDVA.

The above-described method did not allow us to obtain an L-FDVA-derivatized 4-oxo-L-proline. This is plausibly due to the chemical instability of 4-oxo-L-proline in the alkaline condition required for the reaction of derivatization (5).

### Products of 4-oxo-L-proline metabolism in mammalian cell cultures

For the verification of BDH2 expression, lysates of African green monkey COS-7 cells (Cell Lines Service, Eppelheim, Germany) and human HEK293T cells (ECACC via Sigma-Aldrich, Poznan, Poland) were analyzed by western blotting, employing a polyclonal rabbit antibody against BDH2 (PA5-44760, Invitrogen, USA) and a horseradish peroxidase-conjugated goat anti-rabbit IgG antibody (AS09 602, Agrisera), as described previously (25).

To obtain products of 4-oxo-L-proline or *cis*-4-hydroxy-L-proline metabolism in intact COS-7 and HEK293T cells for mass spectrometry analysis, COS-7 or HEK293T cells were plated in 6-well dishes (9.5 cm^2^) at a cell density of 0.35 × 10^6^ or 0.60 × 10^6^ cells/well, respectively, in Dulbecco’s minimal essential medium (DMEM) supplemented with 100 units/ml penicillin, 100 g/ml streptomycin, and 10% (v/v) fetal bovine serum and grown in a humidified incubator under a 95% air and 5% CO_2_ atmosphere at 37°C.

Twenty-four hours after seeding the cells, the culture medium was changed to a fresh one (2 ml) and supplemented with PBS (control), 1 mM 4-oxo-L-proline, or 0.3 mM *cis*-4-hydroxy-L-proline. Next, the cells were incubated for 0, 24, 48 or 72h before transferring the medium (1 ml) to 100 µl of 35% (w/v) HClO_4_. Precipitated protein was separated by centrifugation (13,000 × g for 10 min). After neutralization of the supernatant with 3 M KOH/3 M KHCO_3_, the salts were removed by centrifugation (13,000 × g for 10 min); the clear supernatant was subjected for derivatization with L-FDVA to determine the products of 4-oxo-L-proline and *cis*-4-hydroxy-L-proline metabolism as described above, whereas the consumption of 4-oxo-L-proline was measured spectrophotometrically with the use of recombinant BDH2.

### MTT cell viability assay

COS-7 or HEK293T cells were seeded in 12-well dishes (3.8 cm^2^) at a cell density of 60 × 10^3^ in 1 ml DMEM supplemented with 100 units/ml penicillin, 100 g/ml streptomycin, and 10% (v/v) fetal bovine serum and grown in a humidified incubator under a 95% air and 5% CO_2_ atmosphere at 37°C.

Twenty-four hours after seeding the cells, the culture medium was changed to a fresh one (1 ml) and supplemented with either PBS (control) or 1 mM 4-oxo-L-proline in PBS, and cultured further for 0, 24, 48 or 72 h. To determine the cell viability, 100 µl of MTT (5mg/ml in PBS) was added to each well and the cells were incubated for an additional 30 min in the CO_2_ incubator. After the removal of the culture medium, the intracellular crystals of MTT-formazan were completely solubilized in 1 ml of isopropanol:0.1 M HCl (90:10, v/v). The resulting purple solution was centrifuged (13,000 × g for 10 min) to remove cell debris, and its absorbance was measured spectrophotometrically at λ = 570 nm (formazan) and 690 nm (reference) (26).

### Analytical methods

Protein concentration was determined spectrophotometrically according to Bradford (27) using bovine γ-globulin as a standard. When appropriate, the His_6_-tagged recombinant proteins were detected by western blot analysis, employing a mouse primary monoclonal antibody against His_6_-tag (MA1-4806, Invitrogen, USA) and a horseradish peroxidase-conjugated goat anti-mouse antibody (A2554, Sigma-Aldrich, USA), as described previously (25). All western-blotting analyses employed chemiluminescence and signals acquisition with Amersham Hyperfilm ECL, with the pattern of the prestained protein ladder being copied from the blotting membrane onto the film using a set of felt-tip pens.

### Calculations

V_max_, K_M,_ and k_cat_ for reductase activities of the studied enzymes were calculated with Origin 2020 software (OriginLab, USA) using nonlinear regression. All data are presented as mean ± S.E.

## Data availability

All data are contained within the article.

## Funding and additional information

This work was supported by DSM 501-D114-86-0117600-20 and DSM 501-D114-01-1140100 from the Polish Ministry of Science and High Education and partially by Opus-14 grant from the National Science Centre, Poland (2017/27/B/NZ1/00161). It was carried out with the use of the CePT infrastructure financed by the European Union European Regional Development Fund within the Operational Program “Innovative Economy” for 2007–2013.

## References

1. Semsary, S., Crnovčić, I., Driller, R., Vater, J., Loll, B., and Keller, U. (2018) Ketonization of Proline Residues in the Peptide Chains of Actinomycins by a 4-Oxoproline Synthase. Chembiochem: a European journal of chemical biology 19, 706–715

2. Figueroa M.E., Abdel-Wahab O., Lu C., Ward P.S., Patel J., Shih A., Li Y., Bhagwat N., Vasanthakumar A., Fernandez H.F., Tallman M.S., Sun Z., Wolniak K., Peeters J.K., Liu W., Choe S.E., Fantin V.R., Paietta E., Löwenberg B., Licht J.D., Godley L.A., Delwel R., Valk P.J., Thompson C.B., Levine R.L., Melnick A. (2010) Leukemic IDH1 and IDH2 mutations result in a hypermethylation phenotype, disrupt TET2 function, and impair hematopoietic differentiation. Cancer cell 18, 553–567

3. den Ouden H., Pellis L., Rutten G.E.H.M., Geerars-van Vonderen I.K., Rubingh C.M., van Ommen B., van Erk M.J., Beulens J.W.J. (2016) Metabolomic biomarkers for personalised glucose lowering drugs treatment in type 2 diabetes. Metabolomics 12, 27

4. Mitoma C., Smith T.E., Dacosta F.M., Udenfriend S., Patchett A.A., Witkop B. (1959) Studies on 4-keto-L-proline. Science 129, 95–96

5. Smith T.E., Mitoma C. (1962) Partial Purification and Some Properties of 4-Ketoproline Reductase. The Journal of Biological Chemistry 237, 1177–1180

6. Guo K., Lukacik P., Papagrigoriou E., Meier M., Lee H.W., Adamski J., Oppermann U. 2006. Characterization of Human DHRS6, an Orphan Short Chain Dehydrogenase/Reductase Enzyme. A novel, cytosolic type 2 R-β-hydroxybutyrate dehydrogenase. The Journal of Biological Chemistry 281: 10291–10297

7. Devireddy, L. R., Hart, D. O., Goetz, D. H., and Green, M. R. (2010) A mammalian siderophore synthesized by an enzyme with a bacterial homolog involved in enterobactin production. Cell 141, 1006–1017

8. Langrock, T., García-Villar, N., and Hoffmann, R. (2007) Analysis of hydroxyproline isomers and hydroxylysine by reversed-phase HPLC and mass spectrometry. Journal of Chromatography. B, Analytical technologies in the biomedical and life sciences 847, 282–288

9. Rosenbloom, J., and Prockop, D. J. (1971) Incorporation of cis-hydroxyproline into protocollagen and collagen. Collagen containing cis-hydroxyproline in place of proline and trans-hydroxyproline is not extruded at a normal rate. The Journal of Biological Chemistry 246, 1549–1555

10. Sturm, D., Maletzki, C., Braun, D., and Emmrich, J. (2010) cis-Hydroxyproline-mediated pancreatic carcinoma growth inhibition in mice. International Journal of Colorectal Disease 25, 921–929

11. Yoo, J. S., Sakamoto, T., Spee, C., Kimura, H., Harris, M. S., Hinton, D. R., Kay, E. P., and Ryan, S. J. (1997) cis-Hydroxyproline inhibits proliferation, collagen synthesis, attachment, and migration of cultured bovine retinal pigment epithelial cells. Investigative Ophthalmology & Visual Science 38, 520–528

12. Kanehisa, M., Furumichi, M., Tanabe, M., Sato, Y., and Morishima, K. (2017) KEGG: new perspectives on genomes, pathways, diseases and drugs. Nucleic Acids Research 45, D353–D361

13. Bach, T. M., and Takagi, H. (2013) Properties, metabolisms, and applications of (L)-proline analogues. Applied Microbiology and Biotechnology 97, 6623–6634

14. Persson, B., Kallberg, Y., Bray, J. E., Bruford, E., Dellaporta, S. L., Favia, A. D., Duarte, R. G., Jörnvall, H., Kavanagh, K. L., Kedishvili, N., Kisiela, M., Maser, E., Mindnich, R., Orchard, S., Penning, T. M., Thornton, J. M., Adamski, J., and Oppermann, U. (2009) The SDR (short-chain dehydrogenase/reductase and related enzymes) nomenclature initiative. Chemico-Biological Interactions 178, 94–98

15. Kwiatkowski, S., Seliga, A. K., Vertommen, D., Terreri, M., Ishikawa, T., Grabowska, I., Tiebe, M., Teleman, A. A., Jagielski, A. K., Veiga-da-Cunha, M., and Drozak, J. (2018) SETD3 protein is the actin-specific histidine N-methyltransferase. eLife 7, e37921

16. Liu, Z., Ciocea, A., and Devireddy, L. (2014) Endogenous siderophore 2,5-dihydroxybenzoic acid deficiency promotes anemia and splenic iron overload in mice. Molecular and Cellular Biology 34, 2533–2546

17. Correnti, C., Richardson, V., Sia, A. K., Bandaranayake, A. D., Ruiz, M., Suryo Rahmanto, Y., Kovačević, Ž., Clifton, M. C., Holmes, M. A., Kaiser, B. K., Barasch, J., Raymond, K. N., Richardson, D. R., and Strong, R. K. (2012) Siderocalin/Lcn2/NGAL/24p3 does not drive apoptosis through gentisic acid mediated iron withdrawal in hematopoietic cell lines. PloS One 7, e43696

18. Walejko, J. M., Kim, S., Goel, R., Handberg, E. M., Richards, E. M., Pepine, C. J., and Raizada, M. K. (2018) Gut microbiota and serum metabolite differences in African Americans and White Americans with high blood pressure. International Journal of Cardiology 271, 336–339

19. Geiger, R., Rieckmann, J. C., Wolf, T., Basso, C., Feng, Y., Fuhrer, T., Kogadeeva, M., Picotti, P., Meissner, F., Mann, M., Zamboni, N., Sallusto, F., and Lanzavecchia, A. (2016). L-Arginine Modulates T Cell Metabolism and Enhances Survival and Anti-tumor Activity. Cell 167, 829–842

20. Pan, L., Yu, J., Mi, Z., Mo, L., Jin, H., Yao, C., Ren, D., and Menghe, B. (2018) A Metabolomics Approach Uncovers Differences between Traditional and Commercial Dairy Products in Buryatia (Russian Federation). Molecules (Basel, Switzerland) 23, 735

21. Cantó, C., Menzies, K. J., and Auwerx, J. (2015) NAD(+) Metabolism and the Control of Energy Homeostasis: A Balancing Act between Mitochondria and the Nucleus. Cell Metabolism 22, 31–53

22. Mitsubuchi, H., Nakamura, K., Matsumoto, S., and Endo, F. (2008). Inborn errors of proline metabolism. The Journal of Nutrition 138, 2016S–2020S

23. Summitt, C. B., Johnson, L. C., Jönsson, T. J., Parsonage, D., Holmes, R. P., and Lowther, W. T. (2015) Proline dehydrogenase 2 (PRODH2) is a hydroxyproline dehydrogenase (HYPDH) and molecular target for treating primary hyperoxaluria. The Biochemical Journal 466, 273–281

24. Shevchenko A., Tomas H., Havlis J., Olsen J.V., Mann M. (2006) In-gel digestion for mass spectrometric characterization of proteins and proteomes. Nature Protocols 1: 2856–2860

25. Drozak, J., Chrobok, L., Poleszak, O., Jagielski, A. K., and Derlacz, R. (2013) Molecular identification of carnosine N-methyltransferase as chicken histamine N-methyltransferase-like protein (hnmt-like). PloS One 8, e64805

26. Mosmann T. (1983) Rapid colorimetric assay for cellular growth and survival: application to proliferation and cytotoxicity assays. Journal of Immunological Methods 65, 55–63

27. Bradford M.M. (1976) A Rapid and Sensitive Method for the Quantitation of Microgram Quantities of Protein Utilizing the Principle of Protein-Dye Binding. Analytical Biochemistry 72, 248–254

